# Deep Learning for Sorghum Yield Forecasting using Uncrewed Aerial Systems and Lab-Derived Imagery

**DOI:** 10.1101/2025.07.11.663520

**Authors:** Md Abdullah Al Bari, Aliva Bakshi, Jahid Chowdhury Choton, Swaraj Pramanik, Trevor D. Witt, Doina Caragea, Scott Bean, Krishna Jagadish, Terry Felderhoff

## Abstract

The AI revolution, advanced Graphics Processing Units (GPUs), and open-source platforms have enabled Machine Learning (ML) and Deep Learning (DL) algorithms to rapidly and accurately extract phenotypic features from Uncrewed Aerial System (UAS)-derived imagery. Such advancement leads to phenotypic digitization and sorghum yield forecasting. Yield analytics are critical for breeding programs to assess the genetics and breeding potential of genotypes to enhance cultivar development. This trial followed a three-replicated Randomized Complete Block Design (RCBD) with 36 diverse sorghum genotypes in 2023 at Ashland Bottoms, Kansas. The field images were captured 6 meters above using a DJI M300 drone equipped with the P1 sensor at 90°nadir and 45° oblique angles. This research trained YOLO and the Faster R-CNN (Detectron2) models to harness yield attributes from UAS field and lab images. The YOLO models outperformed the Faster R-CNN model in detecting sorghum panicles, achieving a mean average precision at 50% IoU (mAP@0.50) ranging from 0.92 to 0.98, compared to 0.61 to 0.89. Panicle detection from field imagery correlated at 0.86 with ground truth. Lab imagery analyses measured panicle size, seed counts, and seed area with correlation coefficients of 0.71, 0.95, and 0.25, respectively. Three machine learning models: Support Vector Regression (SVR), Decision Tree Regression (DTR), and Random Forest Regression (RFR) are used to predict yield with correlation coefficients of 0.58, 0.76, and 0.70, respectively. We observed that YOLO models are well-suited for extracting yield-attributing traits from images, which are then incorporated into ML regression models to improve yield prediction performance.

## 1 Introduction

The growing global population drives food production increase of 60-70% by 2050 to meet the projected population growth of 56% [1]. Scientists, including plant breeders, work toward achieving food and nutritional security. Sorghum, a staple in arid and semi-arid regions, is crucial for nutrition, feedstock, and bioethanol production. Sorghum, as a low-input, drought-tolerant [2], phenotypically plastic, and genetically diverse crop, offers the potential for yield and trait improvement. Its resilience to low input, water scarcity, and other stresses makes it an increasingly important crop [3, 4]. The renewed interest in sorghum has driven efforts to accelerate the development of cultivars for local and global food security. Researchers focus on enhancing critical yield attributes through genetic dissection, conventional breeding, and extensive yield trials across locations and years. Sorghum yield is the primary trait of interest for breeders, agronomists, and growers; early, precise, and scalable preharvest yield predictions are crucial but traditionally rely on labor-intensive, time-consuming manual harvesting [5].

Plant breeders need accelerated and accurate tools to record plant measurements in extensive trials for efficient cultivar development. Advancements in remote sensing, high-throughput phenotyping (HTP), and Uncrewed Aerial Systems (UAS) offer innovative approaches to plant phenotyping through high-resolution imagery and videos. Such UAS-derived phenotyping has shown a high correlation with ground truth plant observations. However, the abundance of data captured in large breeding programs presents a challenge in processing and deriving meaningful observations for downstream analysis [6]. Image processing method optimization is critical to effectively leveraging the advantages of these vast imagery resources. Deep Learning (DL) has revolutionized Computer Vision (CV), surpassing traditional image processing to crunch massive and complex datasets to detect objects and efficiently extract meaningful features [7, 8] with the advancement and integration of Graphics Processing Units (GPUs).

Deep Learning, a branch of Machine Learning (ML), leverages deep neural networks such as Artificial Neural Networks (ANN), Convolutional Neural Networks (CNN), Recurrent Neural Networks (RNN), and so on to discover intricate structures in large datasets [9]. Deep Learning excels in processing large datasets by automatically learning a variety of filter parameters during model training. Such capabilities make DL particularly suitable [10] for complex tasks like plant phenotyping, where efficient feature extraction from high-dimensional data is essential. With the advancement of computational speed, parallel computing, and overall efficiency, pre-trained open-source DL models/networks are now available after training on extensive datasets [11]. Such open-source models can be fine-tuned to perform specific object detection or relevant computer vision tasks through transfer learning [12]. This approach circumvents the need to train models from scratch, a computationally intensive and time-consuming process.

Efforts to estimate sorghum yield have leveraged images from mobile applications, focusing on panicle counts and the number of grains per panicle [13], as well as drone-based panicle detection pipelines [14]. Building on these approaches, our work for sorghum yield estimation is taking a step forward, by using UAS-captured red, green, blue (RGB) imagery from the field to develop a training dataset, unlike James et al. [14], where manual hand-held machines were used to create a training dataset. We also extracted yield attributes such as panicle numbers, panicle size, grain numbers, and grain size from these images to forecast sorghum yield.

In this research, we used two open-source, model-driven DL platforms known as You Only Look Once (YOLO) [15] and Faster R-CNN/Detectron2 [16] for object detection. The YOLO platform is widely used in agriculture for detecting crop diseases, pests, and mapping weeds [17, 18, 19]. It has also been applied with greater accuracy in detecting sorghum panicles [14, 20]. Detectron2 is a comprehensive platform for object detection, segmentation, and other visual recognition tasks, offering implementation of various state-of-the-art models, including Faster R-CNN. This model integrates a region proposal network and a detection network that share convolutional features to streamline the region proposal process [21, 22]. This integration results in a highly efficient, accurate, and unified framework for object detection [23].

Forecasting complex attributes often requires specialized ML algorithms, as traditional DL methods may fall short due to their extensive data and computational demands. Support Vector Regression (SVR) has proven effective among various regression models. This algorithm applies the Support Vector Machines (SVM) principles and statistical learning theory [24, 25] to minimize prediction errors. We also employed the Decision Tree Regression (DTR) [26] algorithm to build a single tree that recursively partitions the data into subsets based on feature values, minimizing error [27]. Additionally, we used Random Forest Regression (RFR), an ensemble approach that utilizes multiple decision trees trained on diverse subsets of data. This method leverages a randomly sampled subset of features [28], enhancing its ability to effectively predict complex traits such as yield.

Appropriate prediction can significantly assist breeding programs, agronomy, and plant sciences domains, including enhanced decision making [29] by providing high-quality data promptly. Such efforts enable the quick evaluation of extensive breeding entries, shorten breeding cycles, and efficiently allocate resources toward the most promising genotypes [13, 30].

We hypothesized that DL algorithms could effectively extract yield-attributing features from images as inputs for yield forecasting. This research covered a comprehensive workflow from UAV image collection and data processing to deploying DL frameworks to extract yield-attributing features, fitting these features into ML regression models to forecast yield. Initially, we cropped and labeled UAS imagery and applied YOLO and Faster R-CNN models to detect and count sorghum panicles. Subsequently, panicle sizes were extracted using a fine-tuned YOLO model and the LabelMe [31] bounding boxes. In the following experiments, we employed a fine-tuned YOLO model to count panicle seeds and used SAM 2 [32] masking to measure seed area. By integrating these DL-driven yield-attributing traits into machine learning regressions, we successfully predicted yield with improved accuracy, demonstrating the potential of DL in precision agriculture and crop improvement.

## 2 Materials and Methods

The field trial consisted of 36 US Sorghum lines, laid out in a Randomized Complete Block Design (RCBD) with three replicates at Ashland Bottoms, Kansas, with field co-ordinates 39°8*^′^*20.49*^′′^*N, 96°38*^′^*21.54*^′′^*W (39.1384139, −96.6393167) in 2023. The experimental unit followed 18.5-foot-long 4-row plots, with a row spacing of 30 inches and an alley length of 3 feet. The trial was harvested on November 13, 2023.

### 2.1 Image acquisition

We acquired uncrewed aerial systems (UAS) images from September 25 to 27, 2023, from 2 pm to 5 pm. Using a live video feed, we centered the plot and captured images. The imaging was performed using a DJI M300 (DJI, Shenzhen, China) with a ZenmuseP1 sensor equipped with a 35 mm focal length lens (Figure 1).

**Figure 1:**
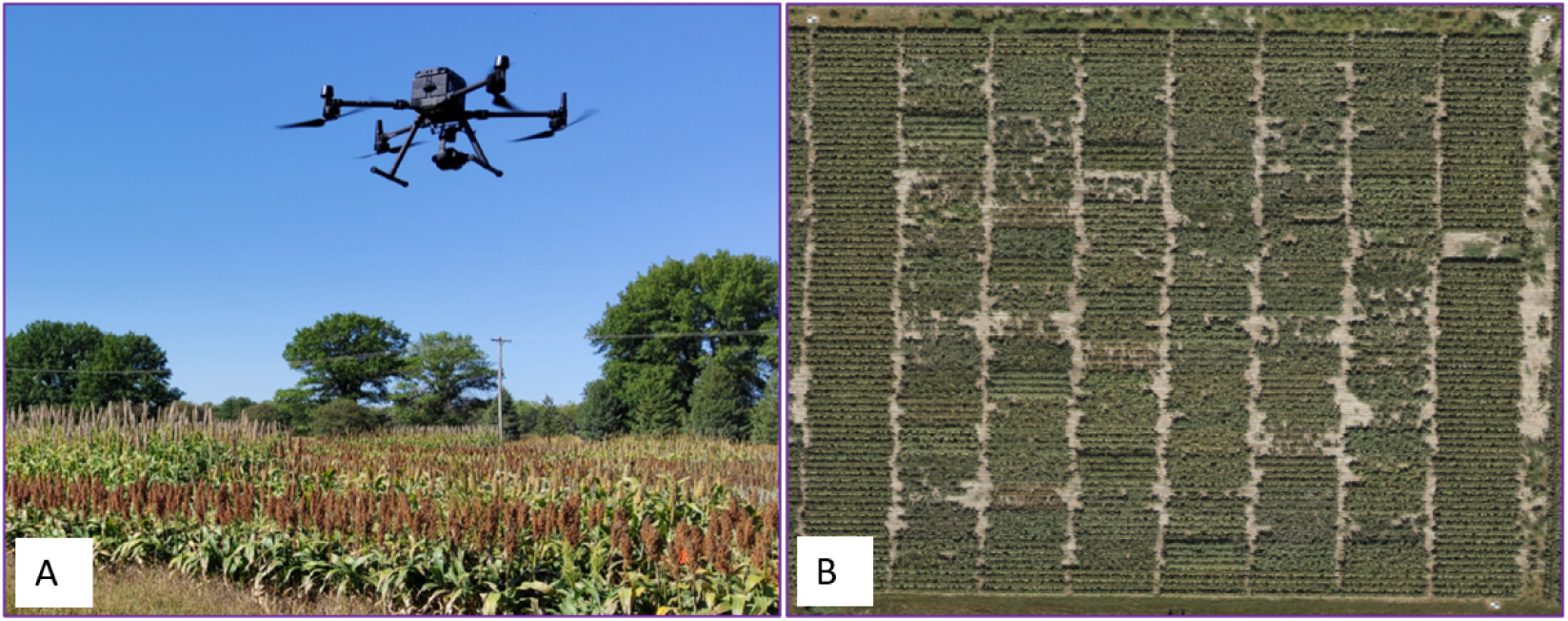
Image capture and view of the Ideotype trial: A. DJI M300 Drone during image acquisition. B. Aerial view of the Ideotype trial at Ashland Bottom, 2023.

During image capture, the flight Above Ground Level (AGL) was 6 m because that was the lowest altitude we could go without severely disturbing the plants (blowing them around), and the shutter speed was 1/500 seconds. The exposure setting was ISO 200 and aperture 5.6 for all flights to achieve a 0.0 exposure. We manually operated this flight. The camera was focused on the center of the plot for each image. For the training dataset, we have images of each line from three angles: Nadir (90° from top), 45° facing north, and 45° facing south. We imaged from three angles because of the shadow variability in each view angle and the uncertainty of which view would potentially perform better. Also, we hypothesize that multiple views can improve the robustness of such detection models. The drone was paused for each image. We captured high-resolution Facing South (FS) 36 images first, followed by 36 Nadir (ND) images, and then 36 Facing North (FN) images. Nadir Ground Sample Distance (GSD) was 0.08 cm/pixel, oblique 0.1 cm/pixel. The white balance was kept sunny. Images were captured with clear skies and low wind (*<* 10 mph). Three images per plot were taken in a single replication (Figure 2).

**Figure 2:**
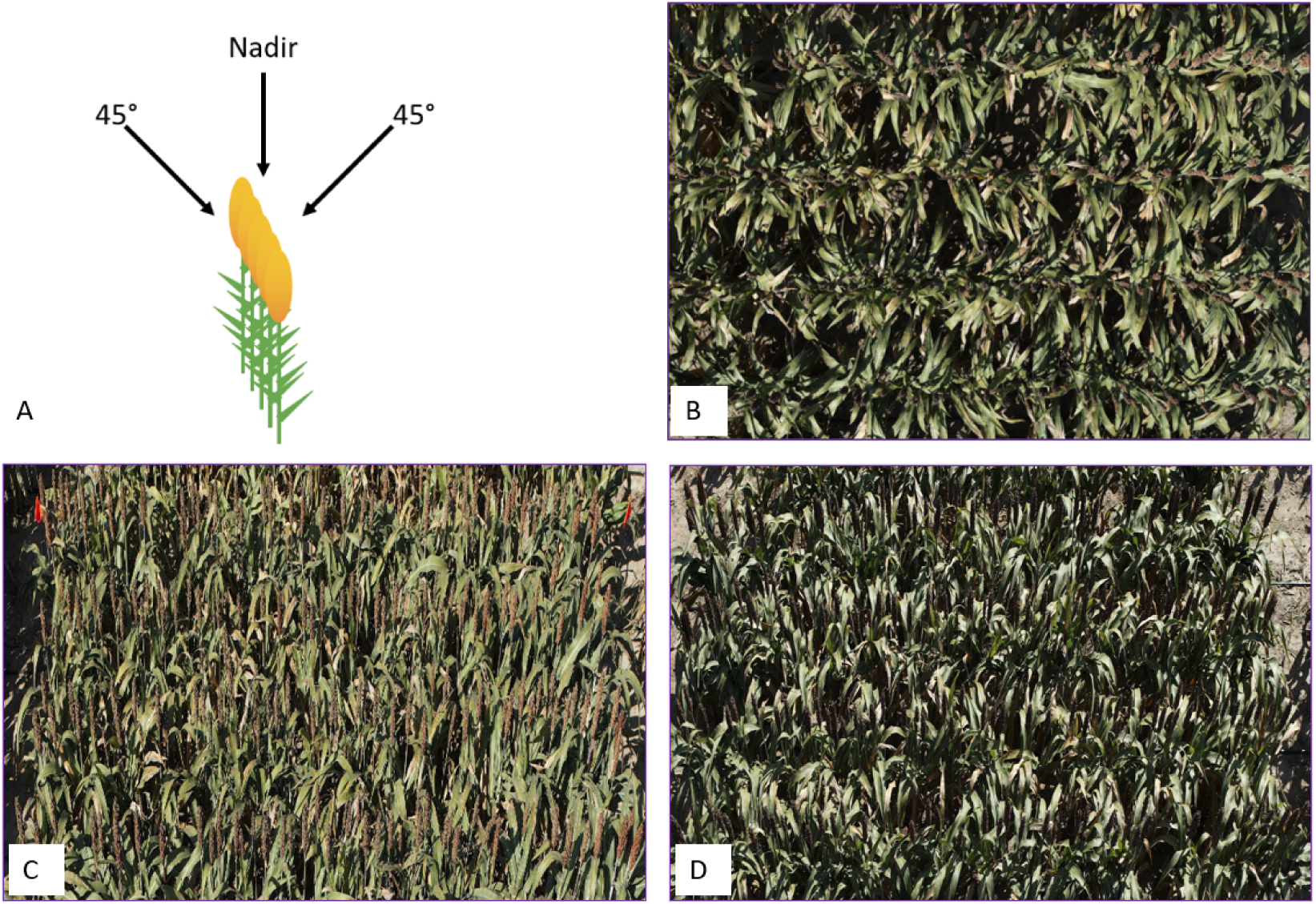
Image captured view angles and perspective of field panicles: A. Cartoon showing three captured angles of field imagery. B. Nadir view of the field panicles. C. Facing north view of the field panicles. D. Facing south view of the field panicles.

### 2.2 Lab data acquisition

We manually harvested four panicles from each plot and set up a stage in a greenhouse to capture images. The stage featured a black cloth backdrop and a measuring scale alongside the panicles. The panicle images were captured from both front and back views. After imaging, the grains were threshed and spread evenly on a black cloth for further imaging. All images were taken using a Canon EOS REBEL SL1 camera with a 37 mm focal length, a resolution of 72 dpi (both horizontal and vertical), and a 24-bit color depth. The distance between the lens and the objects was 38 inches (Figure 3). Machine counts were recorded for each panicle after threshing.

**Figure 3:**
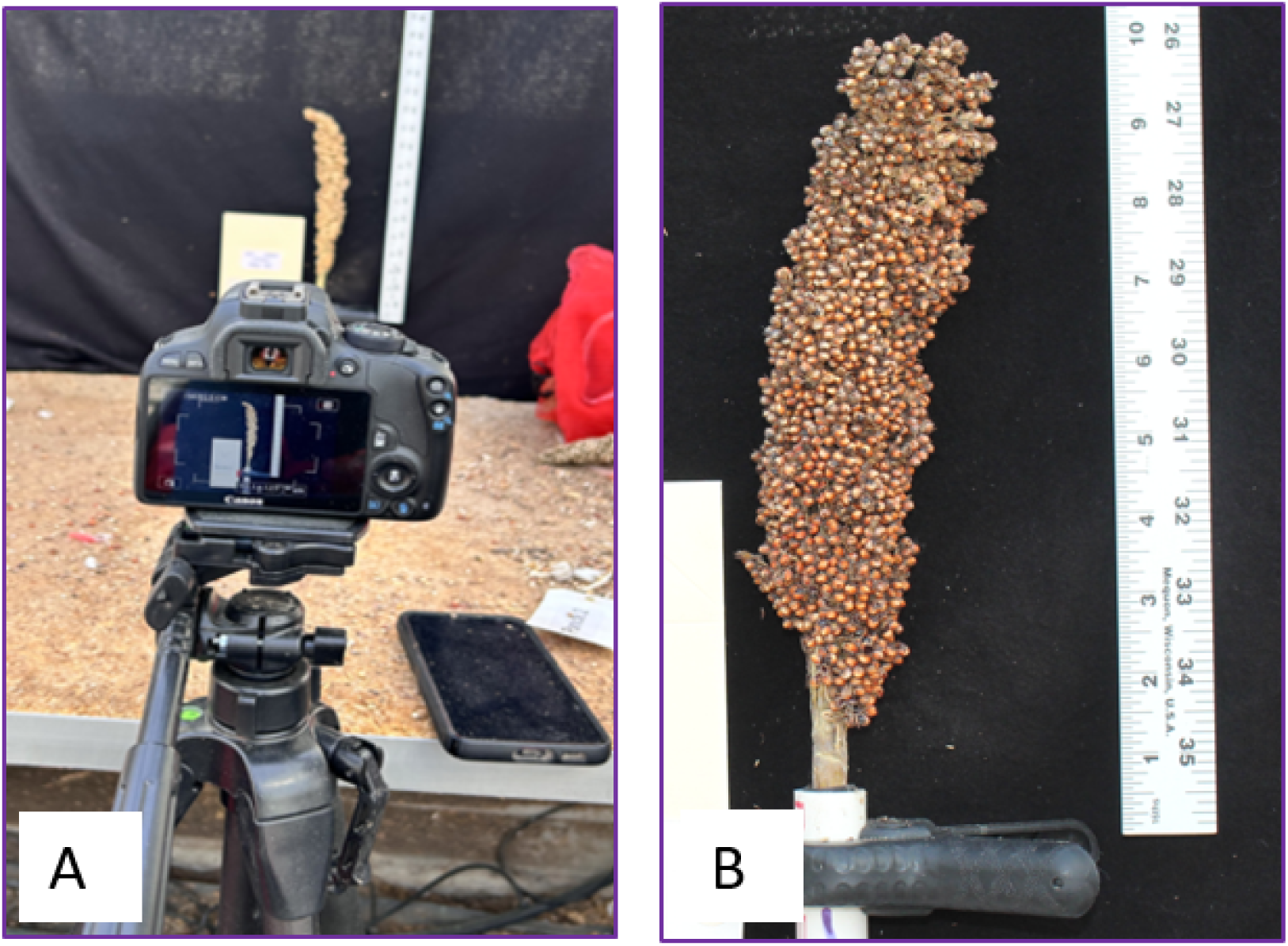
Lab image capture setup for using as ground truth: A. RGB camera to capture harvested panicles. B. Image of captured panicle.

### 2.3 Image cropping and annotation

The captured raw images were large, with an approximate dimension of 8192 pixels (width) x 5460 pixels (height). These images were cropped to 1280 x 1280 pixels and then resized to 640 x 640 pixels (Figure 4).

**Figure 4:**
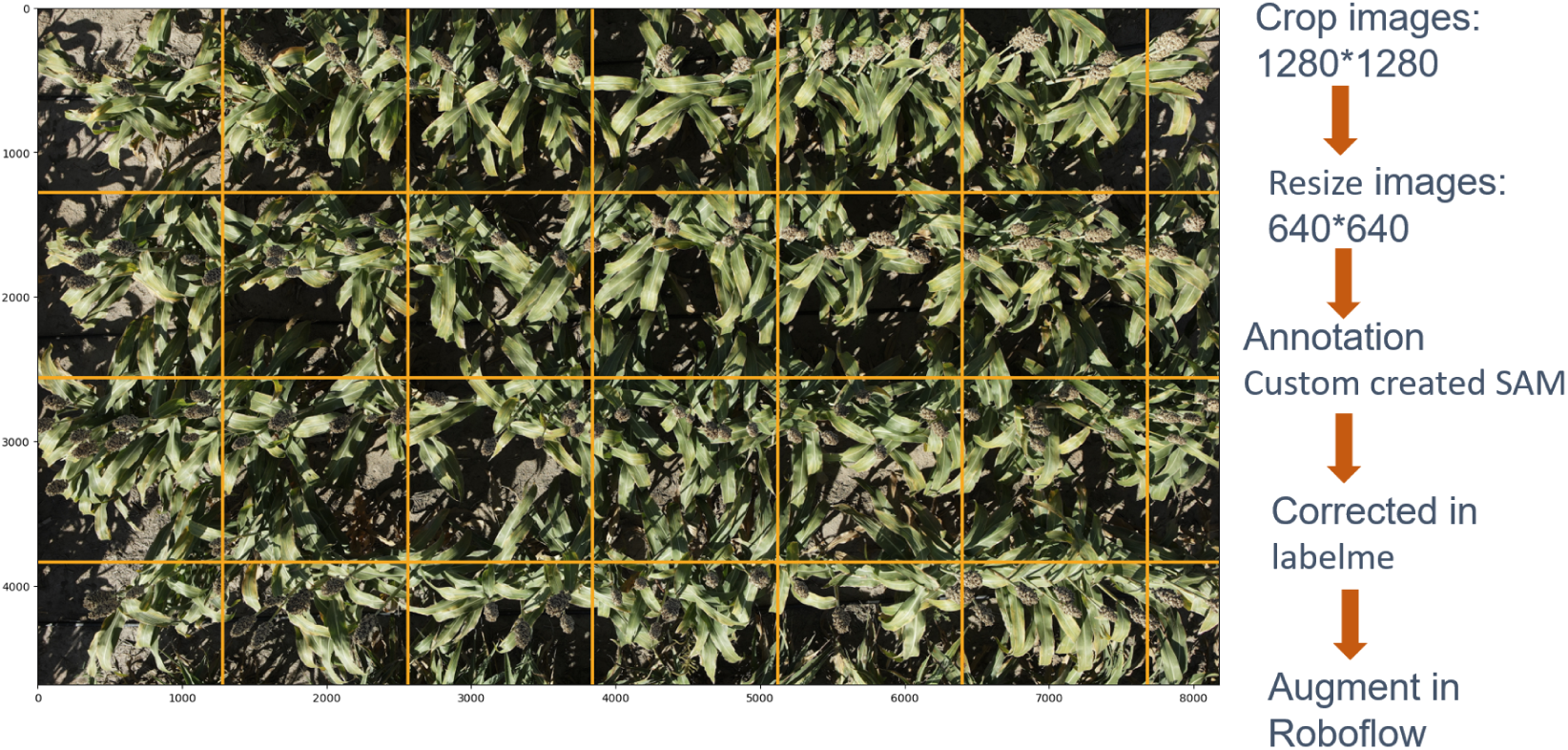
Custom image generation workflow: large image with 1280 x 1280 grids; right panel depicts image cropping at 1280 x 1280 resolution, resizing to 640 x 640, annotation, correction, and augmentation steps.

This allowed the image size to be appropriate for model training in YOLO and Faster R-CNN. We fine-tuned the Segment Anything Model 2 (SAM 2) [32] and generated custom annotation tools based on H-ViT model architecture to segment mask around sorghum panicles quickly (Figure 5). Then, we generated bounding boxes for annotation around the mask. The process resulted in a few incorrect bounding boxes due to overlapping, occlusion, or hiding of panicles.

**Figure 5:**
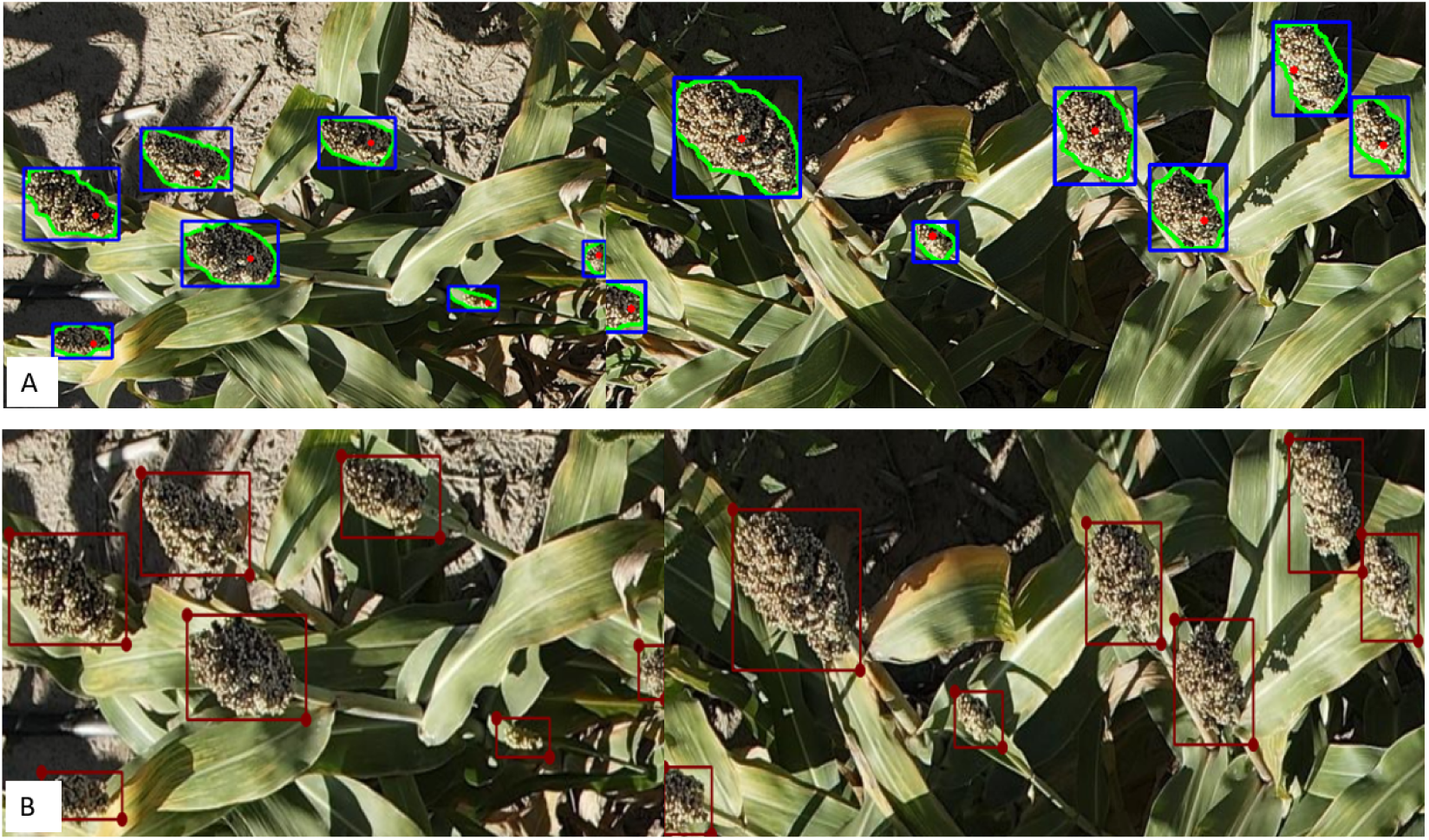
Annotation and Correction: A. Annotated images using fine-tuned Segment Amything Model 2 (SAM 2). B. Correcting annotation using LabelMe.

We used LabelMe [31] to open previously annotated panicle heads (Figure 5) and manually correct these incorrect bounding boxes. This process is faster than manual annotation since a single click generates a segmentation mask, avoiding the need to draw bounding boxes for each panicle.

We developed four types of datasets for panicle detection based on view angles: ND, FN, FS, and All view angles (Table S1). Nadir View started with 303 original images, which, after preprocessing and augmentation, expanded to 841 images, distributed as 807 for training, 17 for validation, and 17 for testing. The comprehensive All view Angles dataset was created by combining images from ND, FN, and FS categories, totaling 1,627 images–1,569 for training, 29 for validation, and 29 for testing, following a 90:5:5 split. We also tried with an 80:10:10 split, with relatively less performance in the prediction; therefore, it was not included. The training, validation, and testing subsets are crucial for developing, fine-tuning, and evaluating the model’s performance. This dataset was designed to enhance the robustness and accuracy of panicle detection. All images are auto-oriented, resized to 640×640 pixels, and converted to grayscale. We included grayscale to prioritize object detection. For further details on other view angles, please refer to Table S1. All preprocessing and augmentation were conducted using Roboflow [33].

### 2.4 DL Models for Object Detection

An object detection model usually consists of a backbone, a neck, and a head. A convolutional neural network (CNN) pre-trained on the ImageNet dataset [34] can form the backbone and generate feature representations. The neck usually consists of more CNN layers and is used for feature enhancement. The head is responsible for identifying objects by predicting bounding boxes around objects and classifying the objects [35]. We used object detection models, such as Faster R-CNN, implemented using the Detectron 2 framework, and three versions of YOLO.

Faster R-CNN is an advanced version of region-based Convolutional Neural Networks (CNNs), initially proposed by Girshick et al. [16] and further developed by Ren et al. [22]. It consists of two neural networks: the Region Proposal Network (RPN) and the Fast R-CNN detector, which work together to identify and classify objects in images. The process starts with an input image passing through a convolutional neural network, such as VGG16, ResNet50, or ResNet101, which produces feature maps containing regions of interest (RoIs) based on pixel groupings. The final output of this model is a highly accurate object detector that predicts bounding boxes, object locations, dimensions, class labels, and confidence scores [36].

YOLO is a one-stage object detector that simultaneously localizes and classifies objects by framing detection as a regression task. YOLO processes full images in a single evaluation step, directly predicting bounding boxes and class probabilities. This unified design enables end-to-end optimization for detection performance, significantly enhancing processing speed. Also, Non-Maximum Suppression (NMS) was applied to refine predictions to eliminate redundant bounding boxes, ensuring the most probable detections [37]. We leveraged YOLO models using the Ultralytics Python library [38].

We deployed YOLOv7, YOLOv8, and YOLOv9 models to compare their strengths in speed, accuracy, and computational efficiency. YOLOv7’s original evaluation showed that it excelled in real-time object detection, outperforming both transformer-based (SWINL Cascade-Mask R-CNN) and convolutional-based (ConvNeXt-XL Cascade-Mask R-CNN) detectors, as well as previous YOLO versions [39]. YOLOv8 was built on YOLOv5 with improved speed, robustness, enhancing detection capabilities, especially for small objects [40], while YOLOv9 introduced Programmable Gradient Information (PGI) and a lightweight Generalized Efficient Layer Aggregation Network (GELAN), which mitigated data loss and further improved accuracy and computational cost, surpassing earlier models like YOLOv7 and the larger YOLOv8-X [41].

### 2.5 Hyperparameter and Training Aspects

In this study, we leveraged multiple advanced object detection frameworks, optimizing each model’s training parameters for enhanced performance. We initiated with Detectron2’s Faster R-CNN, using the R 50 FPN 3x configuration and a base learning rate of 0.007 for 300 iterations, saving checkpoints after 150 iterations. The training configuration processed two images per batch, with Region of Interest (ROI) heads handling 10,240 regions per image and an Intersection over Union (IoU) threshold set at 0.7. The Region Proposal Network was set to manage up to 2,500 proposals post non-maximum suppression.

For YOLOv7, training was conducted using base weights from *yolov*7 *training.pt*, with a batch size of 14 and 100 epochs at an image size of 640 pixels. We aimed for a detection cap of 400 objects per image, employing the *best.pt* weights for inferencing with a low confidence threshold of 0.1 to maximize detection coverage.

YOLOv8 training utilized *yolov*8*s.pt* as base weights, with a configuration designed for 100 epochs, a batch size of 8, and a learning rate of 0.001, maintaining the same object detection and image size parameters as YOLOv7.

YOLOv9 was initialized with *train dual.py* and configured through yolov9-e.yaml. We maintained a constant learning rate of 0.01 across 100 epochs, with momentum set to 0.937 and weight decay set to 0.0005. Detection was performed using *detect dual.py* with *best.pt* to ensure high accuracy during inference. The model training and detection parameters were set to optimize accuracy, efficiency, and computational cost, reflecting the specific operational requirements of each framework.

### 2.6 Measuring Panicle Size

The YOLOv8 model was used to make inferences on the lab-acquired manual images. Panicle length and width were measured from the predicted bounding boxes. We obtained these dimensions by accessing the bounding box data (results[0].boxes.xywh) after prediction. Each bounding box contains the center coordinates (x, y), the width (w), and the height (h). By iterating through the bounding box outputs, we retrieved the width and height of each detected panicle in pixels. As mentioned, we harvested four panicles from each plot and captured images. We trained 324 images, saved the best model, and used the fine-tuned best model to predict inferences in previously untested lab panicles of 108 to estimate panicle sizes. We also utilized the best weights (fine-tuned) saved from the YOLOv8 model training on the FN field dataset to make predictions on the lab-captured panicles after resizing the images to 640 x 640. We hypothesized that the FN view would give better detection in the lab panicles as the FN view had more exposed panicles and better lighting. In this instance, the pixel measurements were not converted into physical units because a reliable conversion factor was not available. Image resizing altered the scale, making the conversion inaccurate. We annotated manually collected laboratory panicle images with rectangular bounding boxes using LabelMe. For each annotation, we extracted the pixel length and width of the panicle using the bounding box recording as (*x*_1_*, y*_1_) and (*x*_2_*, y*_2_) coordinates. The width and height in pixels were computed as the absolute differences *|x*_2_ *− x*_1_*|* and *|y*_2_ *− y*_1_*|*, respectively. The pixel dimensions were then converted into physical panicle length and width measurements by applying the pre-determined pixel counts per inch/cm.

From the field images, we subset panicle images from FN views, as this orientation provided better panicle exposure and sunlight. Using the fine-tuned YOLOv8 model, we detected panicles by projecting bounding boxes around objects. Panicle length and width were extracted in pixels from the bounding box data, similar to the laboratory measurements. These pixel-based measurements were converted into physical dimensions using an appropriate conversion factor. Panicle size is estimated by multiplying the panicle length and width.

### 2.7 Counting Seeds

We collected forward and backward images of four harvested intact panicles from each experimental plot, serving as a paired dataset. Additionally, after threshing each panicle, we imaged the spread grains. To estimate seed count, we applied a fine-tuned seed counting model [42] to detect and count seeds in the forward and backward images of the panicles (Pan Count) (Figure S1 A & B). Since some seeds were occluded and not directly visible, we also employed a counting-by-regression approach (Reg Count), using ResNet50-based regression [43] as suggested by Bakshi et al. [42]. We also estimated seed counts from threshed grain (Spr Count) images (Figure S1 C), where seeds were spread on a backcloth. The machine count of the threshed seeds (Mach Count) was used as the ground truth to evaluate the accuracy of the other methods. We had single rep data across four panicles on 144 observations with very high correlation with the ground truth seed counts (Mach Count). For the second and third reps, we generated the Spr Count by linear regression using the appropriate slope coefficient (0.89) and intercept (273.10), accounting for the standard deviations of the residual of the data in the original 144 observations.

### 2.8 Measuring Seed Area

We developed a graphical user interface (GUI) leveraging the Python library Matplotlib to facilitate interactive seed area calculations. The interface is powered by SAM 2 with H-ViT model architecture [32]. We developed an image viewer class to visualize and interact with images. Users can click on sorghum seeds to initiate object masking, highlighting the seed surface with displayed contours. For each contour, the maximum Euclidean distance, representing the diameter, is calculated between any two points on the contour. This distance is typically measured in pixels.

Sorghum seeds are elliptical in appearance, with one long and one short axis, alternatively, the length and width of the seed. To estimate the length and width of each seed, we utilized the bounding box coordinates generated by the SAM 2 masking. The bounding box dimensions were calculated as follows: the horizontal extent was determined by subtracting the leftmost edge (x1) from the rightmost edge (x2), and the vertical extent was determined by subtracting the bottommost edge (y1) from the topmost edge (y2). Since seed orientation was random, the long axis could align with either the x-axis or y-axis, varying from seed to seed. The greater of these two extents was considered the seed length, while the smaller was classified as the seed width. To convert the pixel distance to standard measuring units, we first convert pixels to millimeters (mm) using the recorded pixel-per-millimeter values from associated metadata. The conversion is defined by:

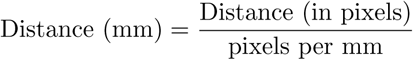

Finally, the seed area in square millimeters (mm^2^) is calculated based on the length and width using the area formula for an ellipse:

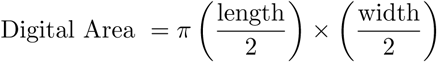

We randomly pointed and clicked 10 to 20 seeds, obtained the seed area, and calculated the means. The graphical interface enables the visualization of seed contours and allows quick and efficient measurement and calculation of seed areas. This platform lays the foundation for capacity building for extensive and rigorous phenotypic evaluation (Figure 6).

**Figure 6:**
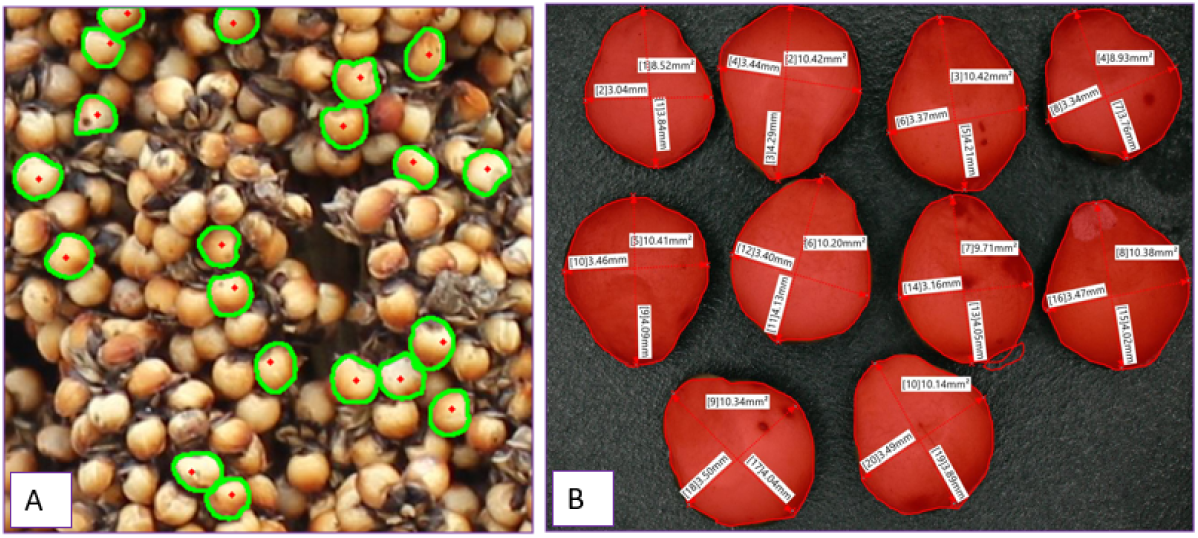
Seed area measurement: A. Digitally obtained seed masks from SAM 2 were used for the digitally obtained seed area. B. Ground truth area of Keyence VX-6000 digital microscope imagery showing the length and the width in mm.

As ground truth, we randomly collected 10 seeds from each of the 108 plots and measured the seed length and width using a Keyence VX-6000 digital microscope (Figure 6). Image analysis tools were used to measure the length and width of the individual kernels (mm). The area of each kernel was measured by tracing the outline of each kernel. Individual kernels were also analyzed using the Single Kernel Characterization System (SKCS) [44] to measure the seed diameter (thickness) (mm) and the seed weight (mg) of the same 10 seeds.

### 2.9 Yield Forecasting

We employed Support Vector Regression (SVR) [24], Decision Tree Regression (DTR) [26], and Random Forest Regression (RFR) [28] to forecast yield (ton/ha) using the Scikit-learn Python library [45]. Yield was the response variable (y), while independent variables (X) included digitally obtained yield attributes: panicle count, lab panicle size (cm^2^), spread seed count, and digital seed area (mm^2^). The dataset was randomly partitioned into training and testing sets using an 80-20 split with a fixed random seed ensuring reproducibility. Scikit-learn does not have any explicit splitting for validation.

The SVR model was initialized with a Radial Basis Function (RBF) kernel–a common choice for nonlinear data. We tuned the SVR parameters with *C* = 1.0 representing the regularization parameter and *epsilon* = 0.1, defining the epsilon-tube within which no penalty is associated in the training loss function with points predicted within a distance epsilon from the actual values. Other model parameters were set to their default values as defined in the scikit-learn implementation of SVR, including: degree = 3, gamma = ‘scale’, coef0 = 0.0, tol = 0.001, C = 1.0, epsilon = 0.1, shrinking = True, cache size = 200, verbose = False, and max iter = −1 [45].

The model fitting process involved multiple iterations over randomly shuffled data splits to ensure robustness and mitigate overfitting. In each iteration, the model was trained on the training set and then used to predict yield on the test set. The mean squared error (MSE) was computed for each prediction to assess the accuracy of the model. This iterative process helped validate the effectiveness of models across different subsets of data. It allowed us to select the iteration with the lowest MSE, representing the model with the best generalization performance on unseen data. These tasks were executed in the testing dataset.

We trained Decision Tree Regression (DTR) and Random Forest Regression (RFR) algorithms using the default hyperparameters in Scikit-learn, following the implementation of SVR mentioned above to predict sorghum yield based on the extracted yield-attributing parameters. The best-performing model was then used to make final predictions on the never-trained 20% test dataset.

### 2.10 Computational Platform Used

The YOLO series models were executed on a Dell Precision 7670 workstation with Python 3.11 development environment under the Linux operating system. This system is equipped with a 12th Generation Intel (R) Core (TM) i7-12850HX processor featuring 24 CPUs and a base clock speed of 2.1GHz, complemented by 65,536 MB of RAM. An NVIDIA RTX A1000 Laptop Graphics Processing Unit (GPU) for enhancing GPU-accelerated tasks. Additionally, computations were performed on an Ubuntu 22.04.4 LTS platform (GNU/Linux 5.15.0-118-generic x86 64), which was outfitted with an NVIDIA RTX A2000 12GB GPU and CUDA version 12.4. We also used Google Colab to train Faster R-CNN models, leveraging a T4 GPU and a Python 3 environment.

### 2.11 Model Assessment and Statistics

In the first experiment, which used four types of imagery data (Table S1), we calculated precision-recall and F1 scores to evaluate the performance of the algorithms on the test and validation datasets [11]. True-positive (TP) and false-positive (FP) detections were determined with an IoU threshold of 0.5 (50%) between the annotated object and predicted bounding boxes. Intersection over Union is the area of overlap between the ground truth bounding boxes and the predicted bounding boxes projected by the trained DL algorithms [46] divided by the total area of the union of the bounding boxes. If the IoU values are greater than or equal to 0.5, they are considered TP, and values less than 0.5 are considered FP [47].

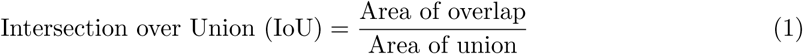

The precision of the model is the proportion of correctly predicted objects (True Positives) to the total number of predictions (TP + FP). The precision values range from 0 to 1. Precision value is reduced with incorrect positive detection, meaning higher FP predictions or fewer TP predictions. A higher precision means lower FP and higher TP predictions. Precision is computed with the equation:

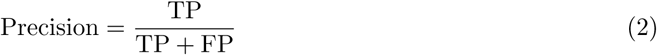

Recall is the fraction of correctly detected objects (TP) out of all ground truth objects (TP + FN), which defines how well a model identifies TP predictions [48]. The higher recall implies better TP predictions, and the value ranged from 0 to 1.

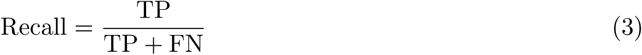

The F1 score is the harmonic mean between precision and recall [49]. In other words, F1 is a function of Precision and Recall. The best score is 1, and the worst score is 0. It was calculated as:

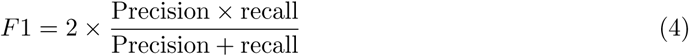

The average precision (AP) for sorghum panicle detection is determined by graphing a precision-recall curve for each test image and computing the area beneath each curve [47]. The AP is then used to find the mean AP (mAP) of the algorithm, which was calculated from an IoU threshold of

0.50 in this study following the equation by Jin et al. [46]:

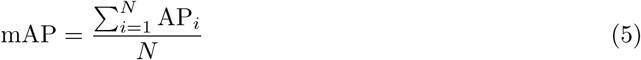

We also assessed AP@0.50-0.95, the mean average precision calculated at multiple IoU thresholds ranging from 0.50 to 0.95. This metric provides a comprehensive measure of detection performance, penalizing models that predict inaccurate bounding boxes. It is more robust than mAP@0.50 since it evaluates precision across different levels of IoU strictness.

In the subsequent experiments, we used fined-tuned model and predicted against ground truth observations. In such instances, we measure the Pearson Correlation Coefficient (r).

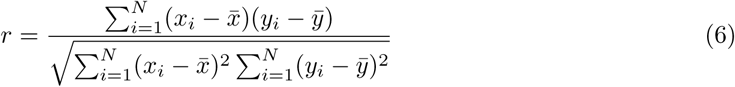

where, *r* is the Pearson Correlation Coefficient, *x_i_* represents model predicted values, *x* is mean of the *x_i_*, *y_i_*refers to actual ground truth observations, and *y* is the mean of the *y_i_* as shown in Equation 6.

Moreover, in all the experiments, we also calculated error metrics, including Mean Absolute Error (MAE), Normalized Error (NE), Mean Squared Error (MSE), and Root Mean Squared Error (RMSE) (Equations 9, 8, 7, and 10) by comparing predictions to the respective ground truth observations to evaluate model performance. Lower values of these error metrics indicate higher model accuracy.

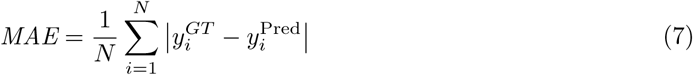

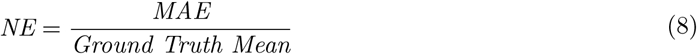

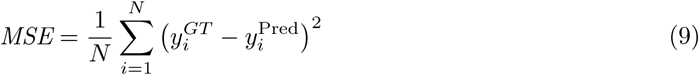

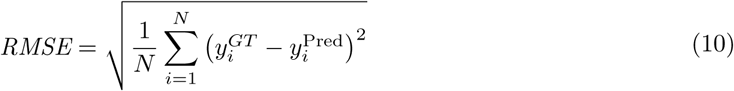

## 3 Results

### 3.1 Field Panicle Detection

We observed varying performance across evaluation metrics using the Faster R-CNN, YOLOv7, YOLOv8, and YOLOv9 models. However, we consistently observed better performance when models were trained using ND and All view angle images to detect panicles. YOLO models performed similarly across versions and image view angles, with AP@0.50-0.95 ranging from 0.64 to 0.90 and AP@0.50 between 0.92 and 0.98 (Table 1). YOLOv7 achieved the highest AP values except for FS, with AP@0.50-0.95 of 0.90 and AP@0.50 of 0.98 when trained on ND view images. YOLOv9 closely followed with an AP@0.50-0.95 of 0.81 and an AP@0.50 of 0.97, while YOLOv8 exhibited slightly lower performance. Though accurate at AP@0.50, Faster R-CNN performed inadequately, especially for FN and FS views. ND view training consistently exhibited improved accuracy, emphasizing suitability for sorghum panicle detection. However, All view images provided a more robust, fine-tuned model after training.

**Table 1:**
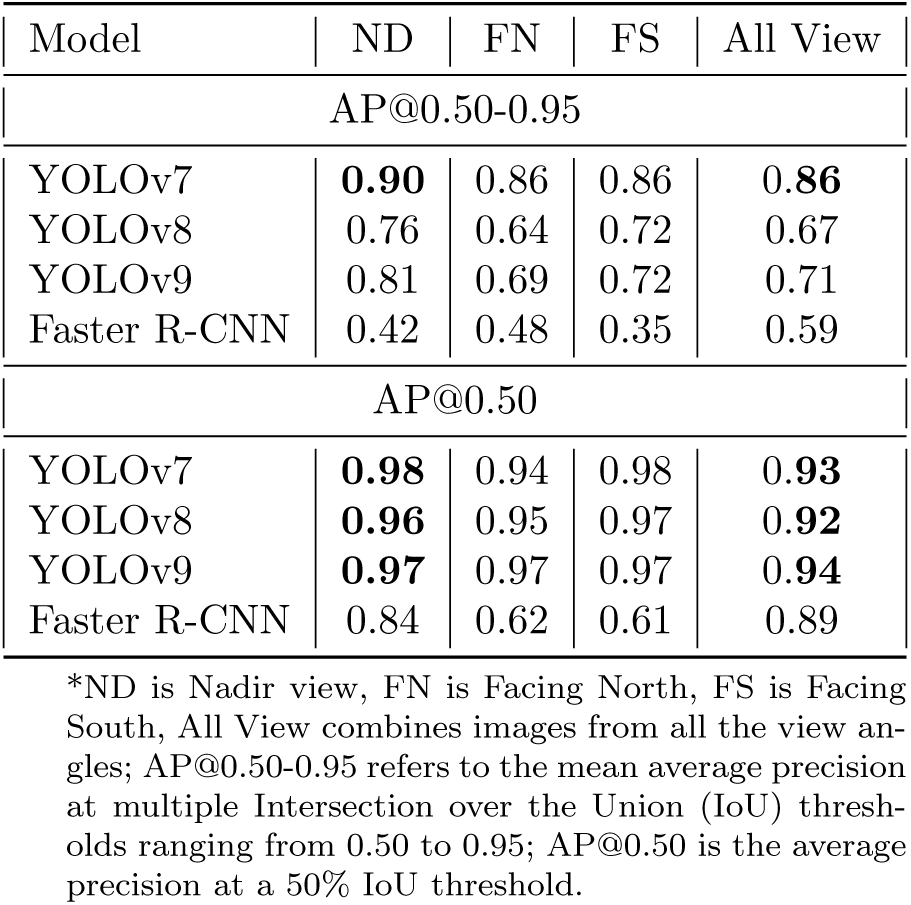
Comparison of Faster R-CNN, YOLOv7, YOLOv8, and YOLOv9 based on AP@0.50-0.95 and AP@0.50 across four view angles.

| Model | ND | FN | FS | All View |
| --- | --- | --- | --- | --- |
| AP@0.50-0.95 |  |  |  |  |
| YOLOv7 | **0.90** | 0.86 | 0.86 | **0.86** |
| YOLOv8 | 0.76 | 0.64 | 0.72 | 0.67 |
| YOLOv9 | 0.81 | 0.69 | 0.72 | 0.71 |
| Faster R-CNN | 0.42 | 0.48 | 0.35 | 0.59 |
| AP@0.50 |  |  |  |  |
| YOLOv7 | **0.98** | 0.94 | 0.98 | **0.93** |
| YOLOv8 | **0.96** | 0.95 | 0.97 | **0.92** |
| YOLOv9 | **0.97** | 0.97 | 0.97 | **0.94** |
| Faster R-CNN | 0.84 | 0.62 | 0.61 | 0.89 |

\*ND is Nadir view, FN is Facing North, FS is Facing South, All View combines images from all the view angles; AP@0.50-0.95 refers to the mean average precision at multiple Intersection over the Union (IoU) thresholds ranging from 0.50 to 0.95; AP@0.50 is the average precision at a 50% IoU threshold.

The F1 score of each YOLO algorithm, as a function of the prediction confidence, is illustrated in Figure 7. The YOLOv7 model achieved the highest F1 score of 0.90 at a confidence threshold of 0.582, while YOLOv8 obtained an F1 score of 0.89 at 0.589. Similarly, YOLOv9 reached an F1 score of 0.90 at a threshold of 0.465. These thresholds represent the optimal balance between precision and recall, ensuring the best detection performance for each model.

**Figure 7:**
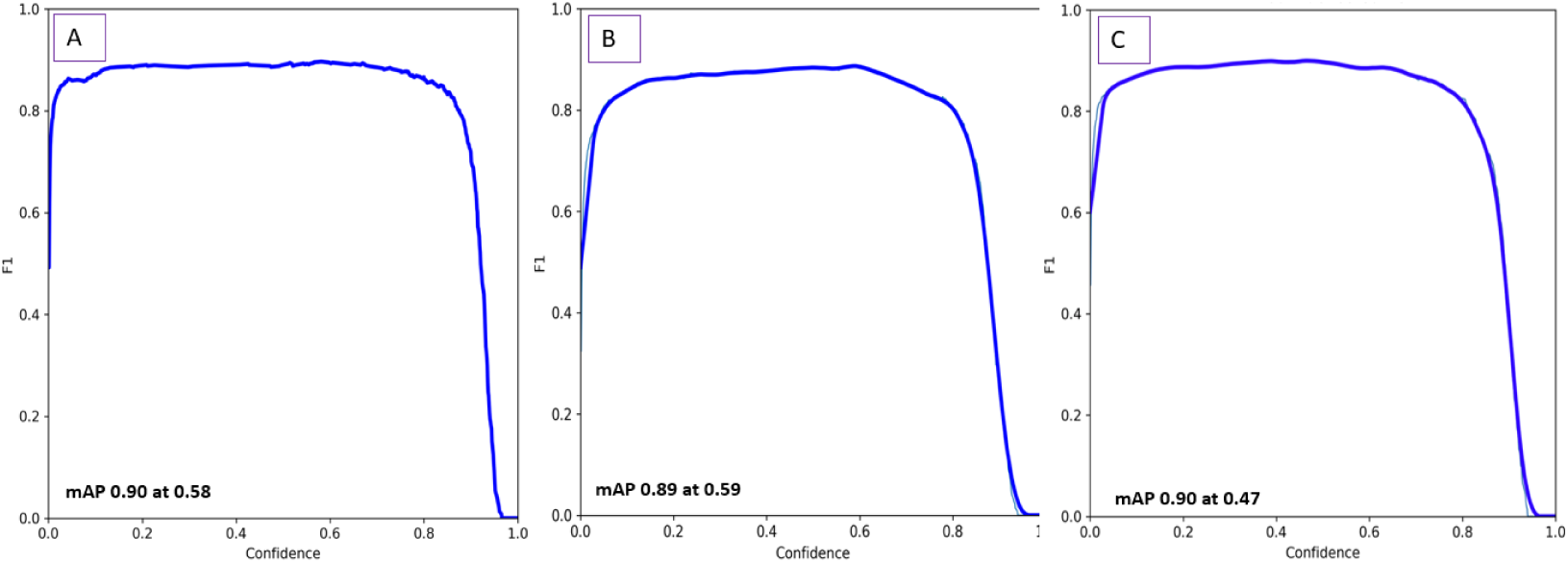
F1 scores training with All view angle images A. YOLOv7. B. YOLOv8. and C. YOLOv9.

The Precision-Recall (PR) curve visually presents the performance of object detection models (Figure 8). YOLOv7, YOLOv8, and YOLOv9 demonstrated greater accuracy in detecting panicles, with mean Average Precision (mAP) scores of 0.93, 0.92, and 0.94, respectively. Since this study focused on single-class object detection, the mAP at an Intersection over Union (IoU) threshold of 0.5 was identical to the class-specific Average Precision (AP) for each algorithm. The results exhibited robust and comparable performance in panicle detection using all three YOLO models per the Precision-Recall (PR) curve.

**Figure 8:**
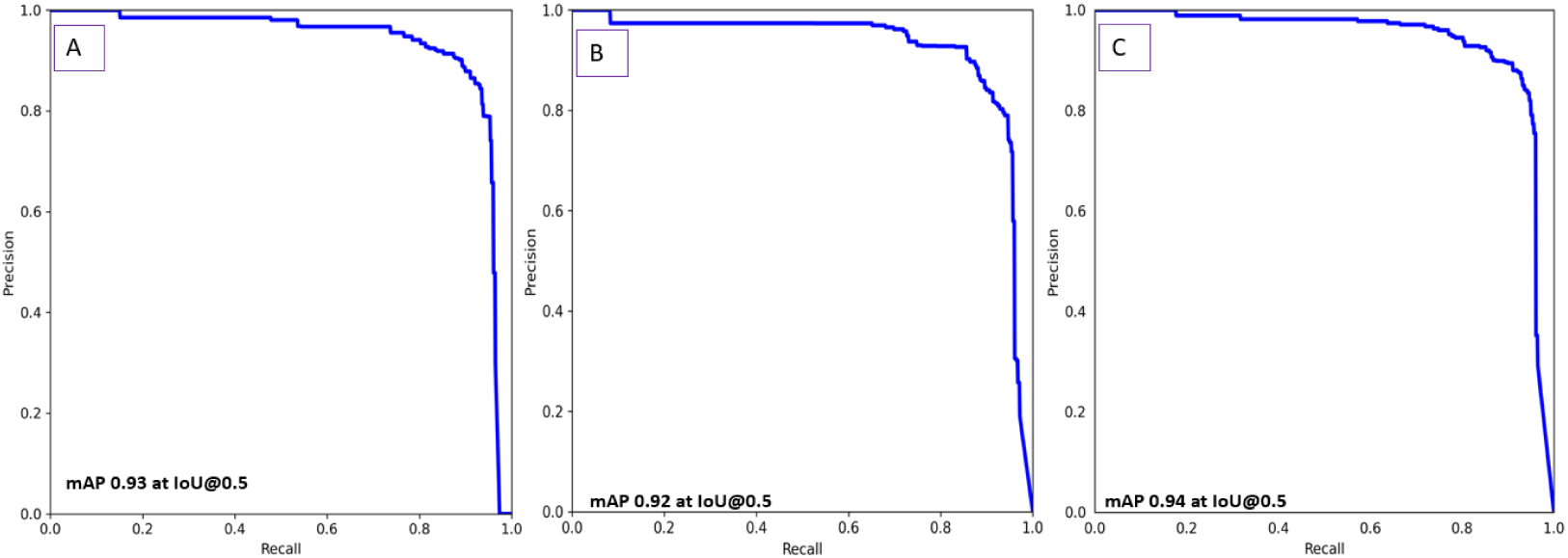
Precision-recall curve while training All view angle images A. YOLOv7 B. YOLOv8 and C. YOLOv9.

The choice of confidence threshold can significantly influence detection performance. Lower confidence thresholds, such as 0.1, tend to yield more detections but are prone to higher rates of false positives, leading to incorrect labeling of panicle heads. In contrast, higher thresholds, such as 0.6, may produce fewer detections with increased precision but at the expense of more false negatives. In this study, confidence thresholds of 0.1 for YOLOv7 and YOLOv9 produced results closely matching ground truth observations, whereas YOLOv8 exhibited higher inference accuracy at a threshold of 0.6.

The detected panicles across view angles and DL algorithms are illustrated in Figure S2. Each YOLO algorithm saved its best-performing weights during training, which we used for inference on large field images. The best-trained model was used to make inferences on the large field images. We found greater accuracy in the ND view images.

Figure 9 illustrates the correlation between the actual ground truth panicle counts and the panicle numbers predicted by the models in the field plots, using fine-tuned models from: A. All view images and B. ND view images. Both the fine-tuned models were used to predict the ND view images.

**Figure 9:**
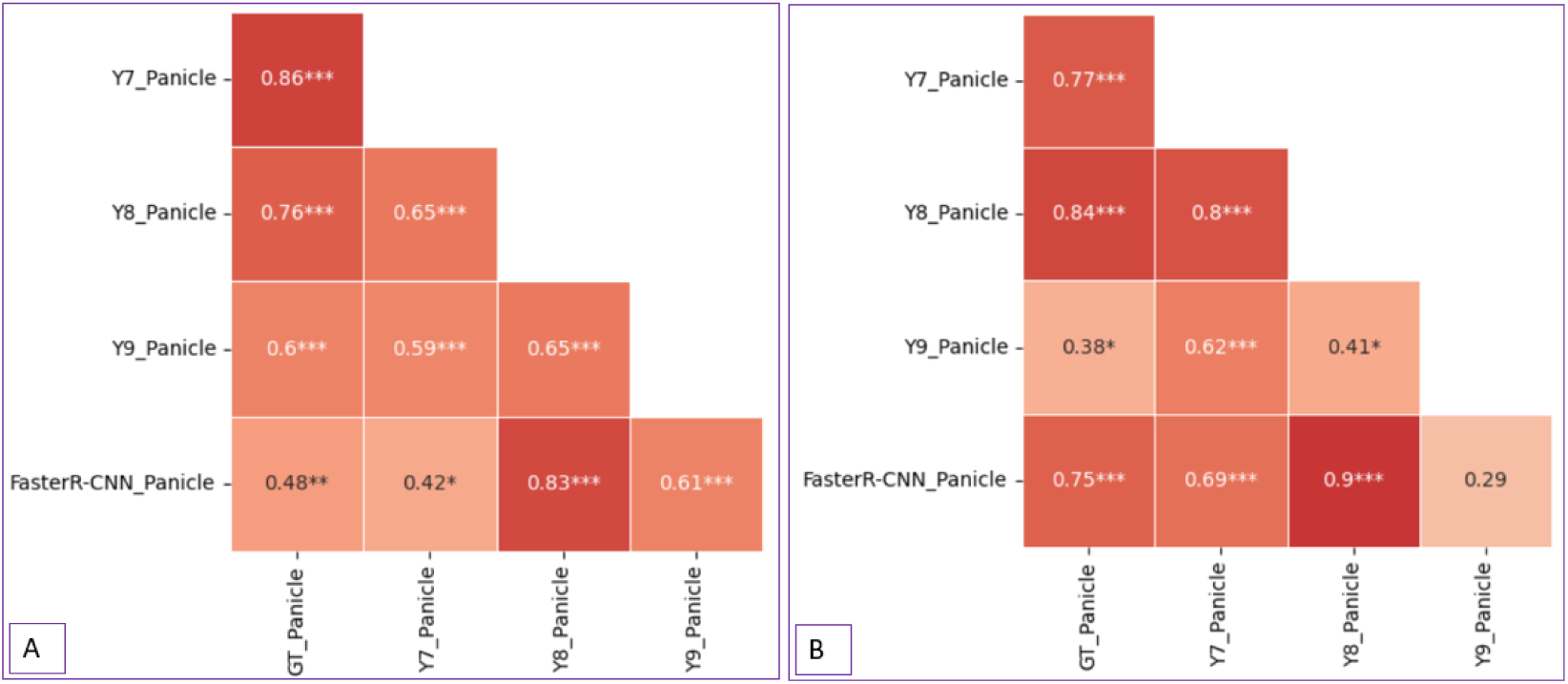
Correlation among ground truth and predicted panicles: A. Model trained with All view angle images. B. Model trained with ND view images. Pearson correlations: * *P <* 0.05, *** *P <* 0.001.

YOLOv7, using the best-trained (fine-tuned) model from All view angle images, predicted very close to the ground truth panicle counts, achieving a correlation of 0.86***, followed by YOLOv8 with a correlation of 0.76***. We obtained slightly varying outcomes when we trained the model with ND view images; YOLOv8 outperformed YOLOv7, achieving a correlation of 0.84***, while YOLOv7 followed a little low with a correlation of 0.77*** (Figure 9 B).

When the best-trained (fine-tuned) model from all-view angle images was used to make inferences on ND-view images, YOLOv8 achieved the lowest MAE (16.42), followed by YOLOv7 (18.97). In contrast, YOLOv9 (48.39) and Faster R-CNN (47.58) performed considerably worse. A similar trend was observed in the NE, where YOLOv8 yielded the lowest NE (0.11), followed by YOLOv7 (0.13), while YOLOv9 (0.32) and Faster R-CNN (0.31) showed substantially higher errors (Table S2).

In the second scenario, where models trained exclusively on ND view images were used for inferences on ND view images, YOLOv8 again outperformed other models with the lowest MAE (14.33), NE (0.09), and RMSE (20.68). YOLOv7 (MAE = 26.89, NE =0.18, and RMSE = 34.83); YOLOv9 (MAE = 72.94, NE =0.48, RMSE = 80.7), and Faster R-CNN (MAE = 23.75, NE = 0.16, RMSE = 28.39) performed moderately (Table S2).

We observed consistent advantages of YOLOv8 across scenarios based on performance; YOLOv7 may also serve as a competitive alternative, while YOLOv9 and Faster R-CNN exhibited lower accuracy and reliability in detecting and counting panicles in the field from UAS imagery.

### 3.2 Panicle Size

One of the reasons we picked the Ultralytics YOLOv8 model is its capability to detect objects with high precision while providing detailed bounding box information. The model generates rectangular bounding boxes for each detected panicles, allowing the extraction of the width and height dimensions. The fine-tuned model trained on the lab imagery was applied to obtain the panicle size. We predicted lab panicle sizes with a correlation co-efficient of 0.59** (Figure S3). The performance matrices included MAE of 46.31 cm^2^, NE of 0.32, and RMSE of 72.33 cm^2^ (Table S3). Additionally, lab panicle dimensions were estimated using LabelMe annotations. The panicle size estimated from LabelMe bounding boxes is strongly associated with the ground truth panicle sizes in square centimeters (cm^2^), with a correlation coefficient of 0.71***. This suggests that LabelMe annotations are reliable for estimating panicle sizes. The further performance matrices included MAE of 38.54 cm^2^, NE of 0.26, and RMSE of 57.52 cm^2^ (Table S3), indicating the accuracy of LabelMe-derived panicle size estimates. Significant correlations were also observed between YOLOv8 predicted panicle size in pixels with the ground truth panicle size in cm^2^ (0.37***) (Figure S3).

### 3.3 Seed Counting

We employed a trained YOLOv8 model [42] to count seeds from the front and back views of the harvested panicles. The counts from both views were summed to estimate the total panicle seed count per panicle. We also counted seeds on spread images after threshing the harvested panicles. A strong correlation was observed between the digitally estimated seed counts and the ground truth measurements. The machine-counted seeds (Mach Count) obtained after harvesting and threshing were used as the ground truth for validation. Among the various seed counting methods evaluated, the spread count demonstrated the strongest association with the machine count (*r* = 0.95***), followed by the panicle count (*r* = 0.73***) and the regression-based count (*r* = 0.56***) (Figure 10).

**Figure 10:**
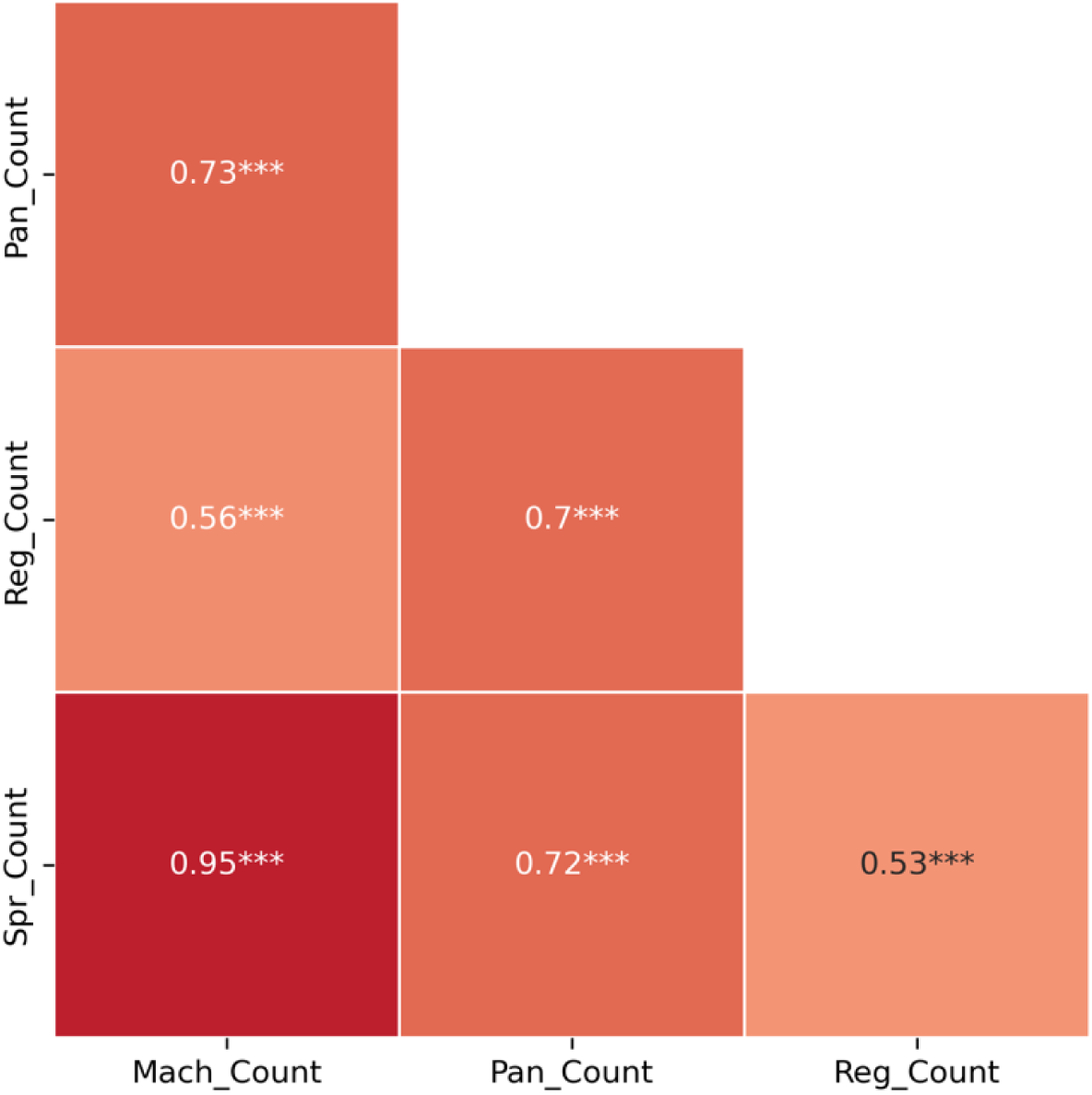
Correlation of machine counted seeds with predicted seeds using a deep learning frame-work. *Mach Count* represents the ground truth machine counts of the harvested and threshed seeds, *Reg Count* is the prediction based on regression count, and *Pan Count* is the sum of panicle counts from the front and back views of a panicle. *** indicates that the Pearson correlation is significant at *P <* 0.001.

The NE ranging from 0.11 to 0.31 is consistent with the correlation trends, and other error statistics are in Table S3.

### 3.4 Seed Area

The seed area was also estimated using a custom-created tool. We calculated the seed area from 10 to 15 random seeds from each genotype. We used another set of 10 randomly selected seeds from the respective genotypes for microscopic seed area (Microscope Area mm2) calculation used as ground truth, SKCS kernel measurement (SKCS Kernel dia mm), and individual seed weight (Kernel weight mg). A moderate but significant correlation existed between the digitally estimated seed area and the ground truth measurements. Among the various seed areas calculated, the digitally obtained area is (*r* = 0.25**), associated with ground truth microscope area estimation(Figure 11).

**Figure 11:**
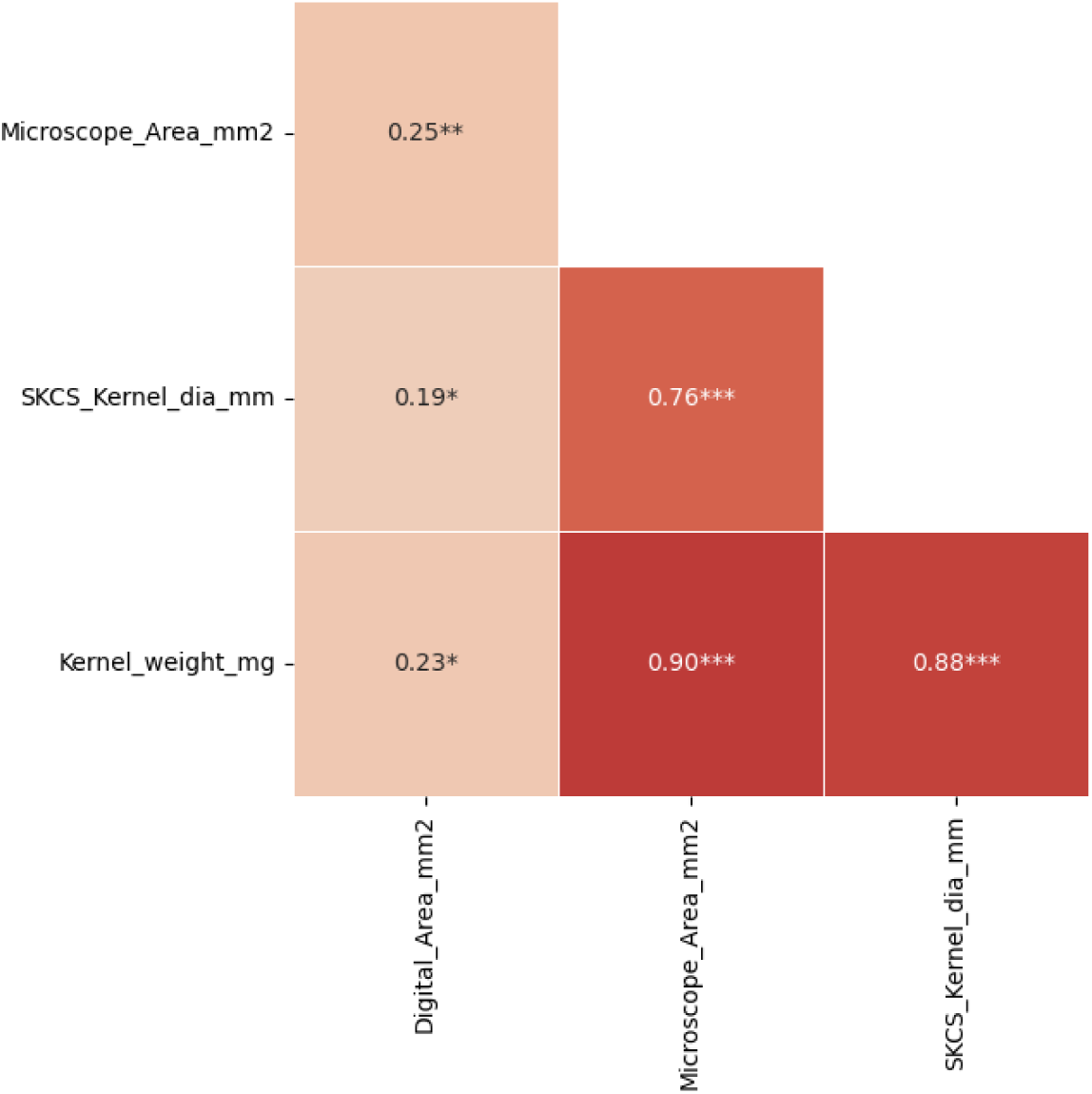
Correlation of digitally obtained seed area with ground truth seed area measurement. *Digital Area mm^2^* represents digitally obtained seed area, *Microscope Area mm^2^* is ground truth area obtained from microscope imagery, *SKCS Kernel dia mm^2^* is Single Kernel ground truth area obtained from microscope imagery, and *Pan Count* is the sum of panicle counts from the front and back views of a panicle. ** indicates that the Pearson correlation is significant at *P <* 0.01.

Digitally obtained seed area (*Digital Area mm^2^*) obtained the least MAE of 2.02 mm^2^, NE of 0.19, and RMSE of 2.71 (Table S3).

### 3.5 Yield Forecasting

We compiled digitally obtained yield attributes and used them as independent variables in the ML regression models to estimate yield, as detailed in the methodology section. The correlation coefficients for SVM, DTR, and RFR on the test dataset were 0.58***, 0.76***, and 0.70***, respectively (Figure 12). Among these models, RFR demonstrated the highest reliability as per error statistics with the lowest mean absolute error (MAE = 0.65 ton/ha), normalized error (NE = 0.19), and root mean square error (RMSE = 0.25 ton/ha). SVR provided MAE = 0.69 ton/ha, NE = 0.21, and RMSE = 0.64 ton/ha, while DTR had MAE = 0.87 ton/ha, NE = 0.24, and RMSE = 1.07 ton/ha (Table S4). Given the average yield of this trial was 3.43 ton/ha, RFR proved to be the most accurate model in our study for predicting yield. However, the highest correlation between the predicted and observed yield was obtained from DRT.

**Figure 12:**
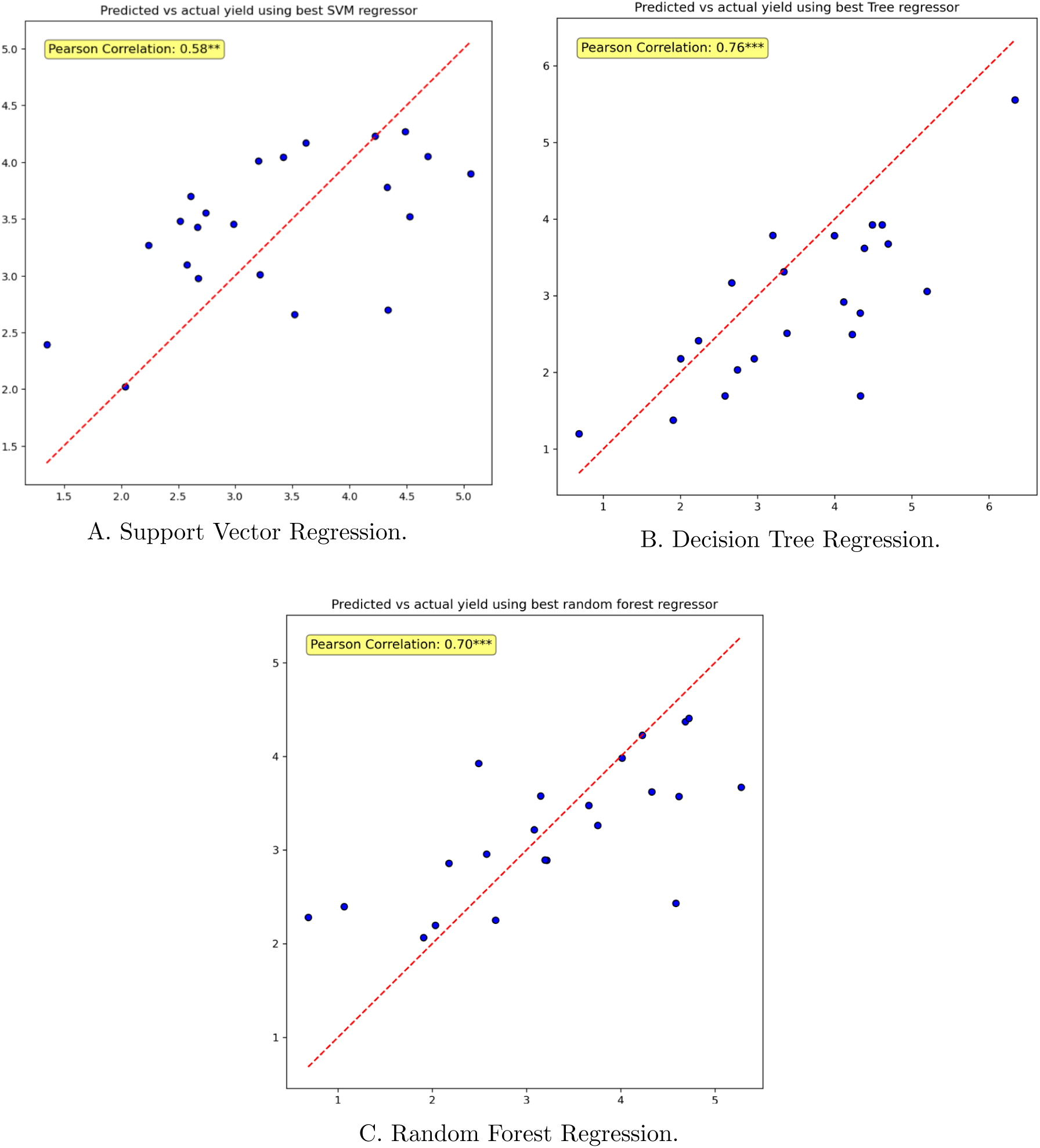
Yield Prediction with Regression Models: A. Support Vector Regression prediction after training and testing on a random dataset, B. Prediction with Decision Tree Regression, C. Prediction with Random Forest Regression. The y-axis is the model predicted yield, and x-axis is the actual yield.

## 4 Discussion

We applied DL learning open-source algorithms in the UAS-derived field images and manually collected lab images to extract yield-attributing traits in the following successive studies:

### 4.1 Field Panicle Detection

The sorghum panicle number or density is a critical determinant of yield [50, 14]. In this study, we assessed the performance of YOLOv7, YOLOv8, YOLOv9, and Faster R-CNN models using various metrics to understand their effectiveness in sorghum panicle detection and localization. The results indicated a clear distinction in performance among the models in detecting sorghum panicles under different training conditions and view angles. While the Detectron2-based Faster R-CNN model showed reasonable performance, YOLO algorithms outperformed specifically when trained with ND and All view angle images.

Among the YOLO series, YOLOv7 and YOLOv8 exhibited reliable object detection across several metrics and view angles when models were evaluated in the test dataset (Table 1). YOLOv7 achieved the highest precision of 0.90 (AP@0.50-0.95) when trained in the ND view images with more distinct panicles, indicating superior feature extraction and localization capabilities in this set of images. The results highlight the potential of YOLO models, particularly YOLOv7, for precision panicle detection in field conditions. While YOLOv8 delivered competitive results, its performance was slightly lower than YOLOv9. Overall, YOLO algorithms reflected high accuracy and robustness in detecting field panicles, showcasing promise for precise object detection-related field phenotyping.

The superior performance of models trained on the ND view images suggested the importance of clearly visible datasets. ND view images provided consistent panicle orientations with minimal occlusions or overlapping as the camera imaged from the top with a 90° angle, enabling more effective feature extraction and learning. In contrast, FN and FS viewed images introduced challenges such as increased occlusions, non-uniform lighting, and shadow effects, complicating object localization and classification. Various panicle orientations in All view datasets addressed these challenges and created a robust training set. Training with such diversified datasets improves the robustness of the model, as evidenced by the consistent performance of multiple YOLO versions (Table 1). This reinforces the importance of varied data to counteract the impact of field-specific imaging conditions. Faster R-CNN detected sorghum panicles less effectively, particularly in FN and FS views, compared to YOLO models. This reduced performance could be attributed to the two-stage approach of the algorithm, which could be less efficient in operating dynamic complexities of field imagery compared to the one-stage architecture of YOLO. Consistent with our findings, other authors reported the sensitivity of Faster R-CNN algorithms to background noise and struggle with detecting small objects [51, 52].

The best weights of the respective fine-tuned models were used to detect panicles in the UAS-derived field imagery of the experimental plots. The strong correlation between ground truth and predicted panicle counts (Figure 9) further signifies the effectiveness of the YOLO models in accurately quantifying panicle numbers from field images. This is critical for high-throughput phenotyping applications, where accurate panicle counts are instrumental for various evaluations. When trained on All view images, YOLOv7 achieved the highest correlation (0.86***) with the ground truth panicle counts, indicating that a diverse dataset enhances the ability of the model to detect field panicles well. YOLOv8 followed closely with a correlation of 0.76***. Although diverse training data improve robustness under varied conditions, it can dilute the ability of the model to optimize for specific scenarios.

Interestingly, YOLOv8 outperformed YOLOv7, achieving a correlation of 0.84*** compared to 0.77*** for YOLOv7 when the trained model weight on ND view images was used to make inferences on ND view images. This suggests that YOLOv8 may better capture specific spatial features or patterns unique to ND view images. These results imply that training data tailored to ND view angle can optimize model performance for that view, highlighting the importance of view-specific training in field-based phenotyping tasks in certain instances. In brief, YOLOv7 and YOLOv8 consistently demonstrated high correlations in All view and ND view angle training scenarios. This suggests their reliability and adaptability in handling variable field image complexities, including variable view angles and backgrounds.

We computed MAE, NE, and RMSE to evaluate the performance of the model in detecting sorghum panicles. Based on these metrics, slightly different results were obtained; YOLOv8 demonstrated superior performance with the lowest MAE, NE, and RMSE values in recognizing field panicles when evaluated with ground truth data in both instances (fine-tuned weight using All view images and ND view images). This suggests that YOLOv8 is better equipped to understand the pattern, generalize, and make accurate predictions, especially for ND view images.

YOLOv7 also showed competitive performance, particularly in All view and ND inference scenarios, as indicated by its MAE, NE, and RMSE. However, its accuracy slightly declined when trained and evaluated exclusively on ND view images (Table S2).

In an 80:10:10 split of train, valid, and test, we obtained comparable but slightly inferior results to the original split. YOLOv7 slightly improved, while YOLOv8 and YOLOv9 got a bit worse. But the results of Faster R-CNN are indeed much worse. The findings show that the YOLO models are more robust than the Faster R-CNN model, whose results change dramatically with variations in the data. We did not include the results of this experiment. Overall, in summary, YOLOv8 consistently demonstrated the highest predictive accuracy across both scenarios, with YOLOv7 showing competitive performance in specific instances. YOLOv9 and Faster R-CNN struggled to achieve comparable accuracy, particularly in more challenging datasets. These findings suggest model specificity and the role of robust training datasets in improving predictive performance.

The performance of YOLOv7, YOLOv8, and YOLOv9 models in detecting panicles has also been evaluated based on F1 scores and Precision-Recall Curve. The F1 scores for YOLOv7, YOLOv8, and YOLOv9 are similar (0.89-0.90). The Precision-Recall curves illustrated the trade-off between precision and recall at different classification thresholds. The Precision-Recall Curve demonstrated mAP scores of 0.93, 0.92, and 0.94 for YOLOv7, YOLOv8, and YOLOv9, respectively, suggesting the comparative strengths of the models in detecting panicles. Guo *et al.* [53] conducted a similar study using a two-step machine-based image processing to detect sorghum panicles from UAS imagery.

Their method achieved a precision of 0.87 and a recall of 0.98 for detecting sorghum heads. Using a geolocation-based plot segmentation method, they reported precision and recall values of 0.82 and 0.98 for panicle detection.

### 4.2 Panicle Size

Sorghum panicle size, defined by length and width, is crucial for crop yield determination as it is a surrogate for the number of seeds. However, accurately measuring panicle size is challenging due to irregular shapes—many panicles are not straight and may exhibit twisted, curved, or tapered forms with narrow tips or bases. Despite these complexities, panicle size remains a meaningful yield-related trait, particularly the area, which represents how many seeds it could hold. This study demonstrates the potential of digital image analysis for estimating sorghum panicle size, showing a strong correlation (r = 0.71***) between manually annotated bounding boxes in lab-captured imagery and ground truth measurements. This high agreement highlights the reliability of human-guided annotation using tools like LabelMe for accurate phenotypic trait estimation. However, when panicle size was predicted using a fine-tuned model trained on the lab imagery, accuracy declined (r = 0.59***), suggesting that while the model performs reasonably well, it does not yet match human annotation precision. This may be due to variability in panicle shape, occlusions, or limitations in the training data. The performance further dropped (r = 0.37***) when a fine-tuned model from FN datasets was used to predict panicle size in pixels compared to physical measurements (cm^2^), indicating challenges in translating pixel-based predictions to real-world units. This could reflect issues in scaling, perspective distortion, or conversion assumptions, especially when working with image-derived data. Panicle size estimation in the lab images was promising but time-demanding, as it is associated with manual imaging and annotations. This finding suggests a need for more automated, yet reliable, tools for high-throughput phenotyping that can reduce labor without significantly compromising accuracy.

The findings suggest that this approach can be extended to panicle size extraction from UAS-derived field imagery. However, field conditions introduced significant panicle size and structure variability, making it challenging to assess predicted panicle dimensions accurately against a reliable ground truth. We extracted panicle length and width from the predictions in a test dataset to evaluate performance. Comparing predicted measurements with ground truth bounding box data, we found that the approach achieved high accuracy.

### 4.3 Seed Numbers

The number of grains in a panicle is a critical trait governing sorghum yield [54, 13]. Accurately counting seeds within a panicle is challenging due to seed occlusion and the three-dimensional structure of the panicle, which limits visibility in two-dimensional (2D) images. The typical imaging approach can cover only the front and back views, omitting the side views. Furthermore, sorghum panicles exhibit significant variation in size, shape, and compactness, concealing internal grains that are not visible externally [55]. These factors contribute to the complexity of achieving reliable seed counts from the sorghum panicles using standard imaging methods.

We implemented a dual-view approach to address some of the limitations in counting seeds from panicles by predicting seed counts from both the front and back views of the imaged panicle using a fine-tuned YOLOv8 model. Summing these predictions resulted in seed counts with a correlation coefficient of 0.73***, compared to ground truth counts. While this method achieved reasonable accuracy for visible seeds, it failed to detect occluded seeds, leading to an underestimation. The observed variation in seed counts was significantly influenced by the shape and compactness of the panicles, which affected seed visibility.

We applied regression-based counting to the panicle images to address concealed seeds. However, contrary to the findings of Bakshi *et al.* [42], our regression-based approach showed a slightly weaker correlation with ground truth counts.

We hypothesized that images of spread seeds from threshed panicles, offering improved seed visibility, would enable the fine-tuned model to recognize seeds more efficiently. The predictions on spread seed images of harvested panicles yielded a strong correlation of 0.95*** with ground truth counts, consistent with our hypothesis. Despite the improved accuracy, this method is time-demanding and requires additional pre-processing, which limits the scalability of large datasets. While developing a non-destructive seed-counting method using field-based UAS imagery remains a desirable goal, the inadequate resolution of current UAS imagery prevented seed counting in this study.

### 4.4 Seed Area

The developed graphical interface, with visualization features, can be used for quick and efficient measurement and calculation of seed dimensions and areas, provided adequate pixel resolution is available. This platform lays the foundation for capacity building for extensive and rigorous phenotypic evaluation of small objects like sorghum seeds.

Sorghum seeds are elliptic in shape, and only the dorsal portion of the kernels is exposed when attached to the panicle. This partial exposure limits the ability of image-based methods to capture the comprehensive appearance of the grains. The lack of proper exposure of kernels in the panicles is a key limiting factor in achieving seed area measurement directly from the panicle.

We obtained a correlation of 0.25*** from the spread images despite extracting the digital seed area from one set of randomly selected seeds while using another set from the same pool for ground truth seed measurement. This platform serves as a foundation for future advancements in phenotypic evaluation of seed areas.

### 4.5 Yield Forecasting

To predict yield (ton/ha), we used digitally obtained yield parameters as independent variables. Machine learning regression algorithms are prevalent in predictions and also for predictive modeling in agriculture. Among the most widely used machine learning algorithms are SVR, DTR, and RFR due to their flexibility and predictive accuracy. Blockeel *et al.* [56], upon studying the role of decision tree regressions over four decades, highlighted the importance of tree-based models to specific datasets for optimal performance. In our research, we applied SVR, DTR, and RFR. RFR provided the highest overall reliability with the lowest normalized error (NE = 0.19), mean absolute error (MAE = 0.65 ton/ha), and root mean square error (RMSE = 0.25 ton/ha). However, DTR achieved the highest correlation between predicted and observed yields (r = 0.76***), suggesting its strength in capturing the structure of the response variable despite having higher error rates than RFR. These findings suggest that both RFR and DTR performed robustly, while SVR showed slightly lower predictive accuracy, possibly due to challenges in kernel selection and tuning with complex, nonlinear datasets [57]. While DTR can be prone to overfitting, especially in the absence of proper pruning or appropriate tree-depth in the decision process [58, 27], its performance in our study indicates effective fitting. RFR, as an ensemble method, mitigates overfitting by averaging multiple decision trees, handles feature interactions more effectively, and performs well with high-dimensional, nonlinear data [28, 59]. Thus, RFR proved to be an accurate and reliable model for yield prediction in this trial, with DTR also showing promise.

In consonant with our findings, Yamparla *et al.* [60] found Random Forest as the best yield predictor with an accuracy of 95% after training from temperature, rainfall, yield, and pesticide data. Savaliya *et al.* [61] studied the predictive performance of yield on weather and soil parameters using K-Mean, Linear Regression, Random Forest, and Support Vector Machine. Savaliya and colleagues reported a hybrid approach of K-Mean merged with Random Forest, which provided yield estimation with an accuracy of 99.77%. Contrary to our findings, Yasaswy *et al.* [62] reported that Innovative Gradient Boosting (98.7%) outperformed Random Forest (86.4%) on a small dataset.

### 4.6 Limitations

The variable shape and posture of field panicles [8], complex background, occlusion of seeds in the panicle [42], and insufficient pixel resolutions in the UAS-derived images were some of the key limitations that hindered the precise extraction of yield attributes. Despite these challenges, we generated substantial datasets and object detection protocols to extract yield-attributing traits with greater accuracy. We could not fly a drone (UAV) lower than 6m to get higher-resolution images because that would have disturbed the sorghum plants. We would need a longer focal lens and/or a higher megapixel sensor to get higher-resolution images. When trained on relatively small datasets, RFR demonstrated greater precision. However, a more robust dataset is required to effectively evaluate the predictive performance of the predictive regression analytics. Enhancing data collection methods with more precise sensors can enable higher-resolution imaging, addressing these constraints. Additionally, camera and sensor technology advancements, standardized viewpoints, optimized lighting conditions, and improved scaling methods can further enhance the accuracy of extracting yield-attributing traits from the collected imagery and improve yield prediction.

## 5 Conclusion

Advancements in AI, DL, and ML hold immense potential in advancing agricultural research [13, 63, 14]. With advanced computational power and various ML algorithms, plant breeders can efficiently extract key yield-related attributes, gain rapid insights into genotype performance, simplify, and improve selection processes. For example, breeders can eliminate a substantial proportion of less promising candidates early in the breeding cycle, such as the bottom 70%, narrowing their focus to thoroughly evaluate top-performing genotypes. Such a targeted approach enhances breeding decisions in identifying suitable genotypes with desirable traits and accelerates progress toward desirable goals [64].

Automated yield prediction is a pressing demand to keep pace with the need to reach breeding goals rapidly. The findings from our panicle detection study highlight the performance of the YOLOv8 and YOLOv7 models over Faster R-CNN in terms of precision and localization accuracy. However, YOLOv8 was deemed more accurate in the large field plot images in localizing panicles. The measured panicle size from the field was difficult due to the lack of proper resolution in capturing the panicle size variability. Still, we obtained greater accuracy between the predicted bounding box and ground truth labeling in a subset of the testing dataset. We obtained greater accuracy in the lab panicle sizes. Seed counting and seed area estimation were also promising using our DL platforms.

Accurate and automated sorghum yield forecasting is cardinal for breeding selections, farm management, and resource allocation. This research aims to revolutionize traditional agricultural practices by integrating DL frameworks with UAS-derived panicle imagery to create an efficient pipeline for extracting yield attributes. This approach promises early yield prediction by leveraging the extracted yield attributes into ML regression algorithms. Integrating SVR, DTR, and RFR, we obtained prediction accuracies of 0.58, 0.76, and 0.70 with the actual yield. The findings demonstrate the potential of DL-extracted features to forecast yield and thus could assist in transforming crop management, breeding, and policy decisions, accelerating high-yielding cultivar development. Our research also presented a pipeline for extracting yield attributes for forecasting sorghum yield by integrating DL frameworks with UAS and manual imagery, contributing to the digitization of sorghum and other breeding and research programs.

## Author Contributions

**Md Abdullah Al Bari**: Conceptualization; methodology, image preparation, analyses, predictive model; writing – original draft, review, editing. **Aliva Bakshi**: Faster R-CNN/Detectron2, YOLO models, seed counting analysis, review, editing. **Jahid Chowdhury Choton**: SAM 2 based annotation tool, YOLO models, panicle count predictions, panicle size, seed area analysis, review, editing. **Swaraj Pramanik**: Faster R-CNN/Detectron2, YOLO models, seed counting analysis, review, editing. **Trevor D. Witt**: UAS image acquisition, review, editing. **Doina Caragea**: Conceptualization, resources, overseeing analyses, review, editing. **Scott Bean**: Ground truth seed area measurement, review, editing. **Krishna Jagadish**: Funding acquisition, resources, review, editing. **Terry Felderhoff**: Conceptualization, resources, overseeing analysis, review, editing.

## Acknowledgments

Dennis Hitz and Matt Davis led hourly help, a crucial part of this research. Thanks to Arthur Selman, Computer Systems Specialist in the Department of Agronomy, for maintaining the Milo server. Mention of trade names or commercial products in this publication is solely for the purpose of providing specific information and does not imply recommendation or endorsement by the United States Department of Agriculture (USDA). USDA is an equal opportunity provider and employer.

## Data Availability

After preliminary acceptance, the datasets, fine-tuned model weights, and codes will be publicly available in the GitHub repository.

## Supplementary Materials

**Figure S1:**
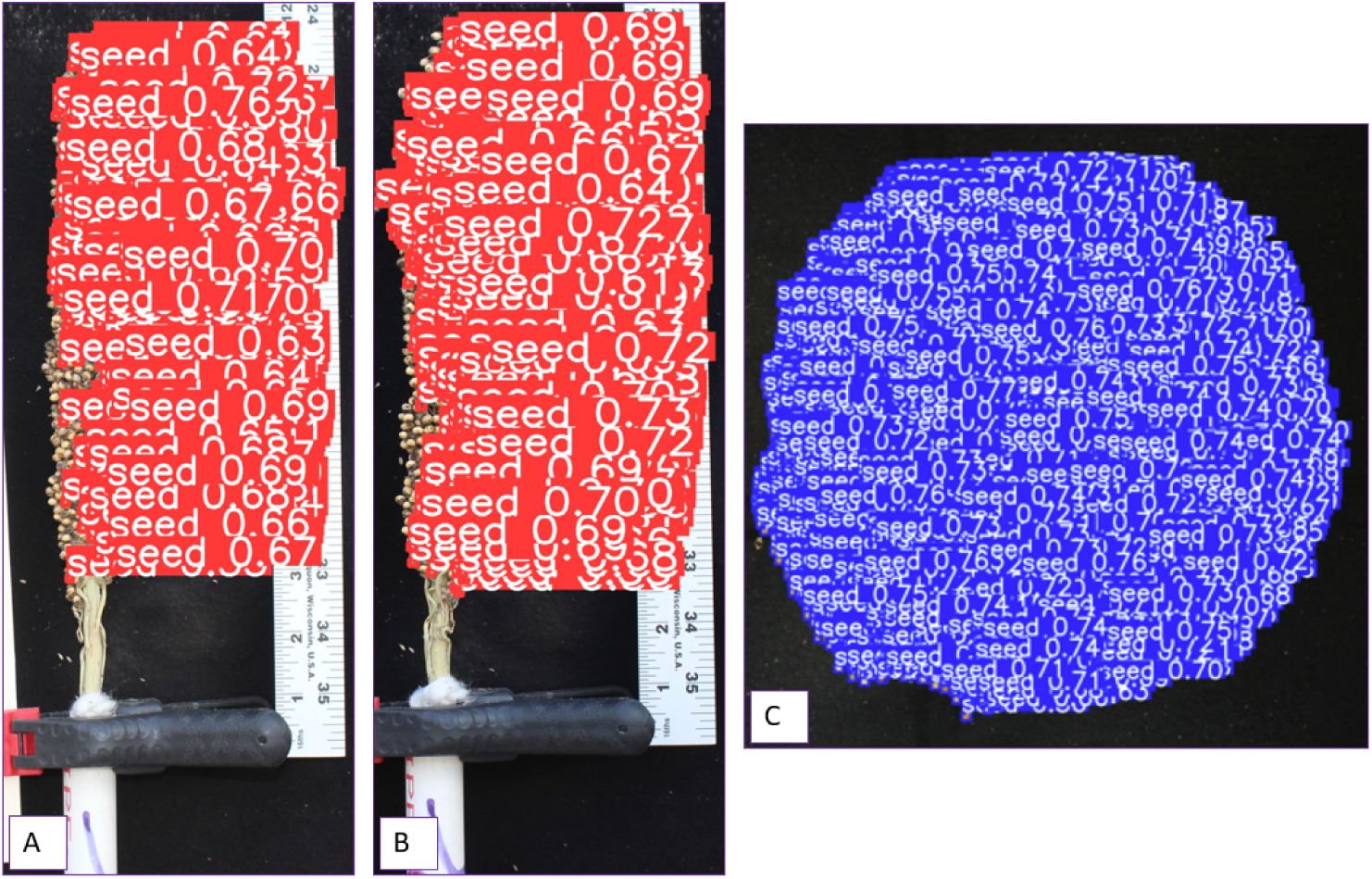
Detected seeds A. Panicle forward view detection B. Panicle backward view detection C. Detection on the spread images

**Figure S2:**
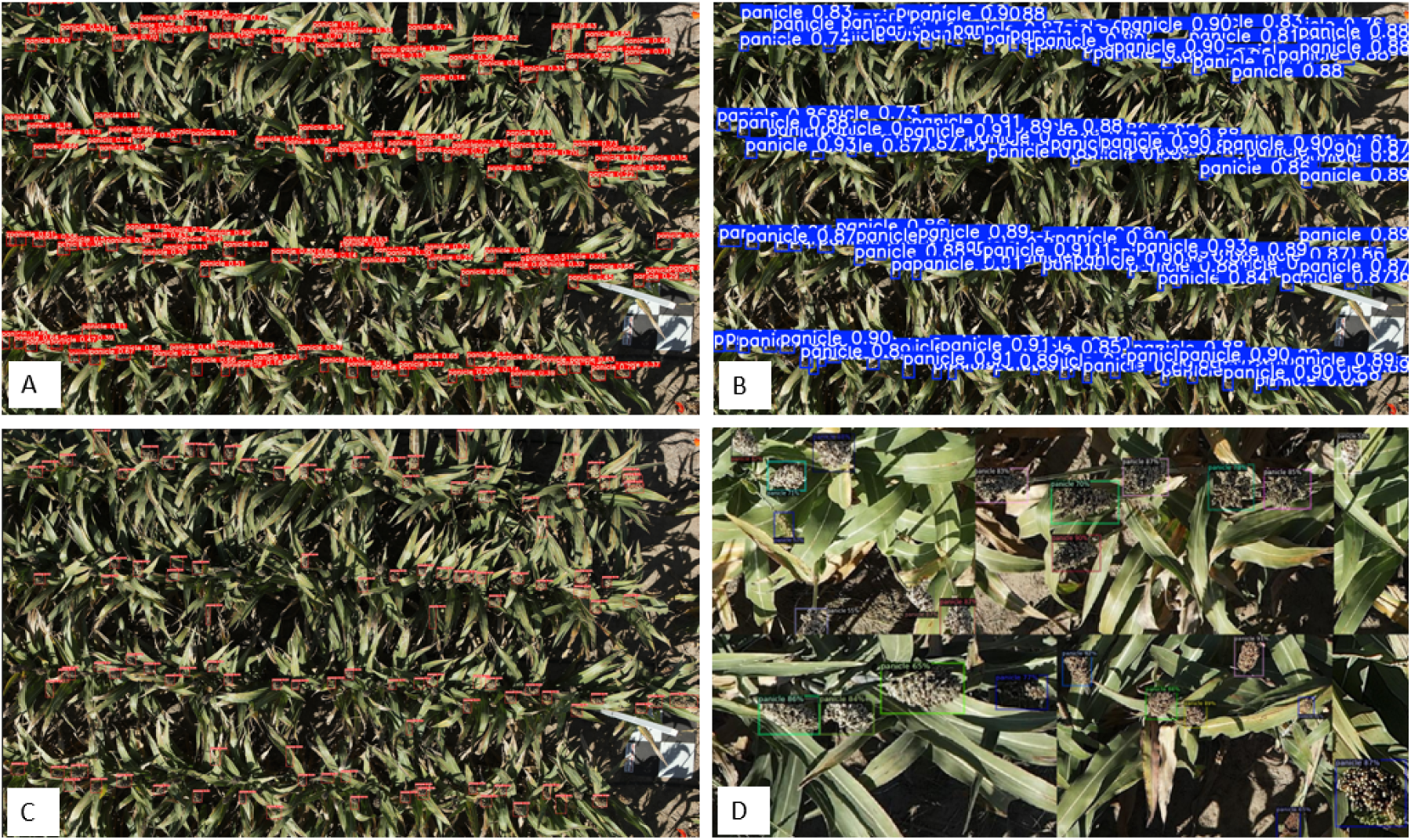
Detected panicles on Nadir view images with A. YOLOv7 B. YOLOv8 C. YOLOv9 and D. Faster R-CNN. We used 640 x 640 sliding windows to detect panicles and summed for the entire image as the model was unable to detect objects in a large image size of approximately 8000 x 5400.

**Figure S3:**
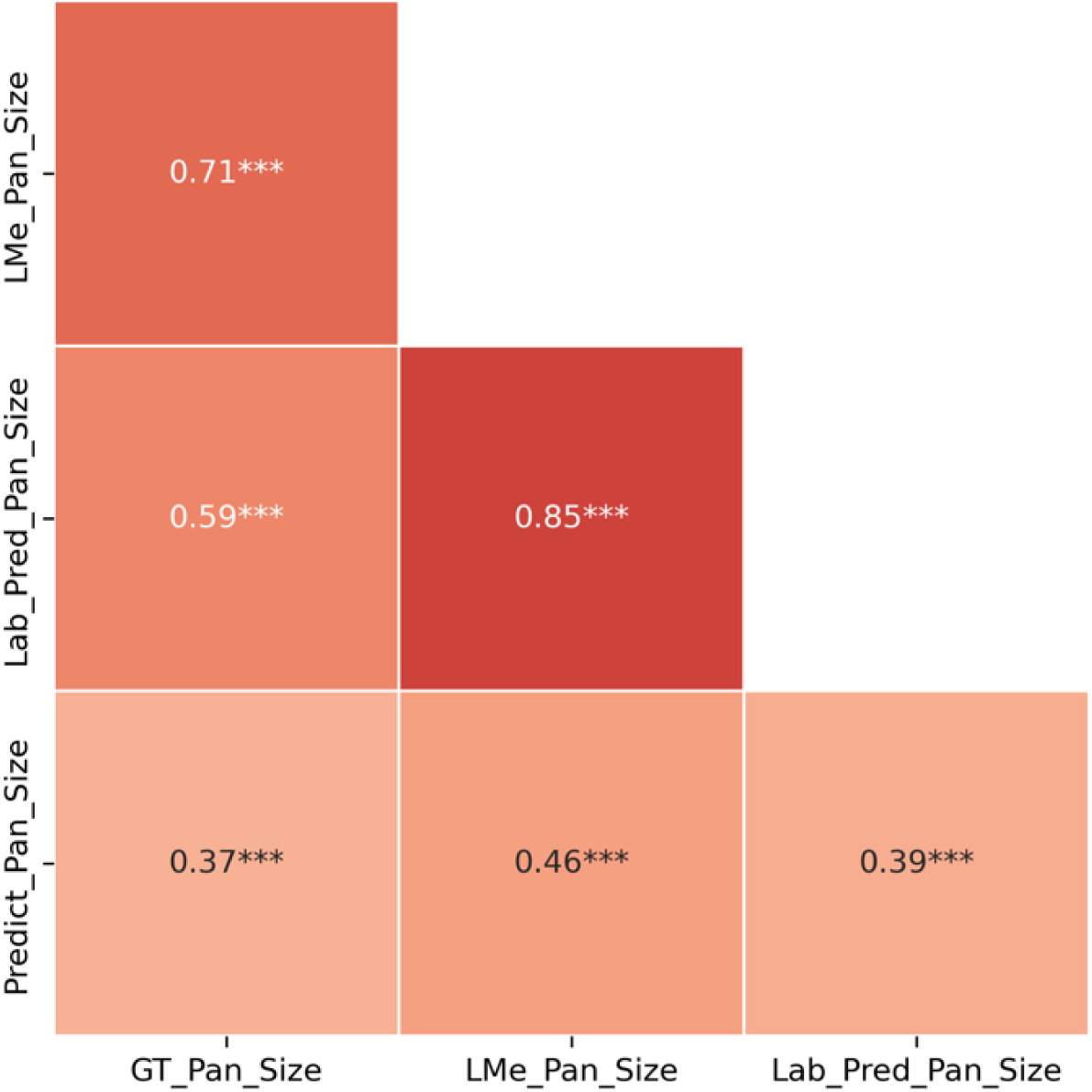
Correlation co-efficient of digitally obtained panicle size vs ground truth panicle size. *GT Pan Size* represents the ground truth panicle size (cm^2^) measured from lab imagery (length *×* width). *LMe Pan Size* refers to the panicle size (cm^2^) extracted from LabelMe-derived bounding boxes, *Lab Pred Pan Size* refers to the panicle size (cm^2^) extracted by predicting bounding boxes from the trained lab imagery, while *Predict Pan Size* denotes the predicted panicle size in pixel^2^ from the fine-tuned trained model. *** indicates that the Pearson correlation is significant at *P <* 0.001

**Table S1:** Generated images for panicle detection from different view angles.

| View Angle | Initial Images | Augmented Images | Training | Validation | Testing |
| --- | --- | --- | --- | --- | --- |
| ND | 303 | 841 | 807 | 17 | 17 |
| FN | 200 | 614 | 594 | 11 | 9 |
| FS | 195 | 547 | 528 | 10 | 9 |
| All Views | 581 | 1627 | 1569 | 29 | 29 |

\* ND: Nadir view, FN: Facing North, FS: Facing South.
\* All Views include combined images from multiple view angles.

**Table S2:** Comparative accuracy of Deep Learning models in detecting panicles using ND view imagery.

| Statistics | YOLOv7 | YOLOv8 | YOLOv9 | Faster R-CNN |
| --- | --- | --- | --- | --- |
| **Trained with All View Angle Images** |  |  |  |  |
| MAE | 18.97 | 16.42 | 48.39 | 47.58 |
| NE | 0.13 | 0.11 | 0.32 | 0.31 |
| MSE | 537 | 659 | 3085 | 3338 |
| RMSE | 23.17 | 25.67 | 55.54 | 57.78 |
| **Trained with ND View Images** |  |  |  |  |
| MAE | 26.89 | 14.33 | 72.94 | 23.75 |
| NE | 0.18 | 0.09 | 0.48 | 0.16 |
| MSE | 1213 | 428 | 6513 | 806 |
| RMSE | 34.83 | 20.68 | 80.70 | 28.39 |

\* DL: Deep Learning, MAE: Mean Absolute Error, MSE: Mean Square Error, NE: Normalized Error, RMSE: Root Mean Square Error.
\* **All View Angle Images:** Images combined from multiple view angles.
\* **ND View Images:** Nadir view (top-down) images.

**Table S3:**
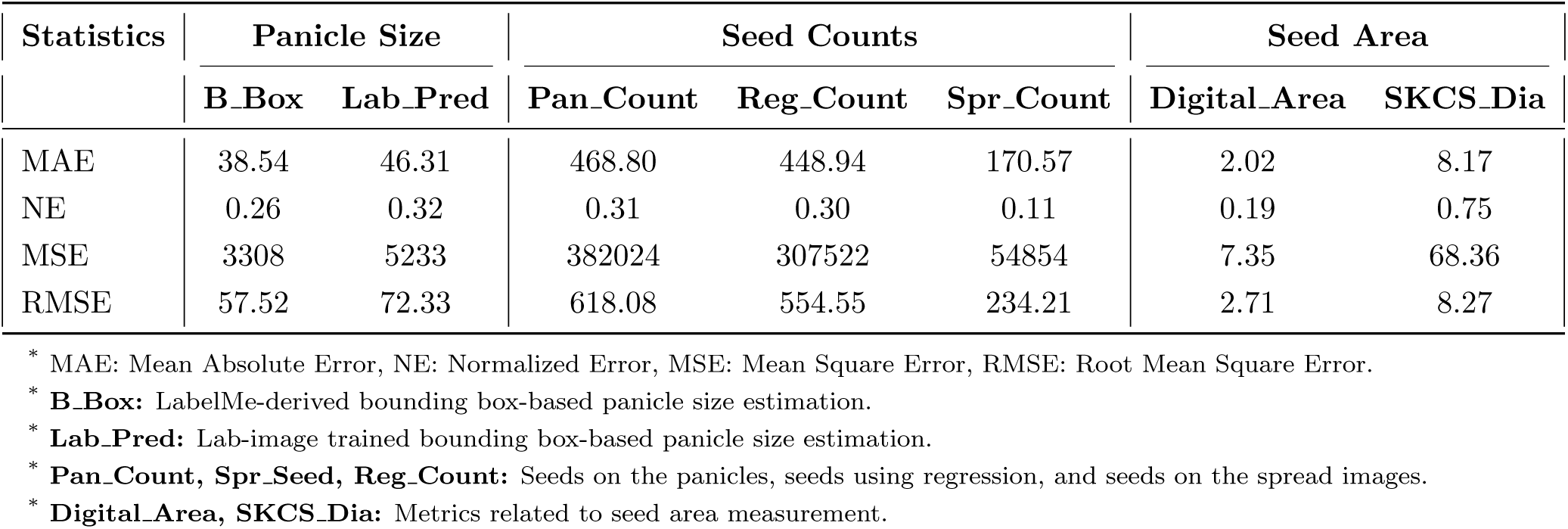
Statistics related to the prediction accuracy compared with the respective ground truth observation.

| Statistics | Panicle Size |  | Seed Counts |  |  | Seed Area |  |
| --- | --- | --- | --- | --- | --- | --- | --- |
|  | B_Box | Lab_Pred | Pan_Count | Reg_Count | Spr_Count | Digital_Area | SKCS_Dia |
| MAE | 38.54 | 46.31 | 468.80 | 448.94 | 170.57 | 2.02 | 8.17 |
| NE | 0.26 | 0.32 | 0.31 | 0.30 | 0.11 | 0.19 | 0.75 |
| MSE | 3308 | 5233 | 382024 | 307522 | 54854 | 7.35 | 68.36 |
| RMSE | 57.52 | 72.33 | 618.08 | 554.55 | 234.21 | 2.71 | 8.27 |

\* MAE: Mean Absolute Error, NE: Normalized Error, MSE: Mean Square Error, RMSE: Root Mean Square Error.
\* **B\_Box**: LabelMe-derived bounding box-based panicle size estimation.
\* **Lab\_Pred**: Lab-image trained bounding box-based panicle size estimation.
\* **Pan\_Count**, **Spr\_Seed**, **Reg\_Count**: Seeds on the panicles, seeds using regression, and seeds on the spread images.
\* **Digital\_Area**, **SKCS\_Dia**: Metrics related to seed area measurement.

**Table S4:** Statistics related to the yield prediction accuracy of ML regression models.

| Statistics | Support Vector Regression | Decision Tree Regression | Random Forest Regression |
| --- | --- | --- | --- |
| MAE | 0.69 | 0.87 | 0.65 |
| NE | 0.21 | 0.24 | 0.19 |
| MSE | 0.64 | 1.15 | 0.77 |
| RMSE | 0.78 | 1.07 | 0.88 |

\* ML: Machine Learning, MAE: Mean Absolute Error, MSE: Mean Square Error, NE: Normalized Error, RMSE: Root Mean Square Error.
\* Models evaluated: Support Vector Regression, Decision Tree Regression, and Random Forest Regression.

